# Genome-wide mapping of mesoscale neuronal RNA organization and condensation

**DOI:** 10.1101/2025.04.19.649570

**Authors:** Lindsay A. Becker, Sofia A. Quinodoz, Troy J. Comi, Ofer Kimchi, David A. Knowles, Clifford P. Brangwynne

## Abstract

Subcellular RNA organization can affect critical cellular functions. However, our understanding of RNA microenvironments, particularly biomolecular condensates, remains limited, largely due to a lack of technologies to comprehensively interrogate mesoscale RNA organization. Here, we adapt Split-Pool Recognition of Interactions by Tag Extension to map micron-scale RNA-RNA spatial proximity genome-wide across cell regions (RNA-SPRITE). Deploying RNA-SPRITE, we find extensive, conserved organization of mature mRNAs, with increased colocalization between mRNAs that share RNA-binding protein (RBP) motifs or encode functionally related proteins. Both effects are especially strong in dendrites and axons, suggesting prevalent mRNA co-regulation. Moreover, mRNAs with less compact folding, lower translation efficiency, and specific RBP motifs are more likely to be in RNA-rich condensates. However, perturbations that broadly dissolve or enhance condensation reveal that RBP motif and encoded protein-mediated colocalizations largely remain intact, independent of condensation. These results demonstrate the power of RNA-SPRITE in revealing critical aspects of RNA’s functional organization.

**In Brief:** Unbiased, genome-wide maps of RNA-RNA mesoscale spatial proximity uncover extensive subcellular organization and its governing principles.

**Highlights:**

- RNA-SPRITE reveals micron-scale RNA colocalization genome-wide across cell regions
- mRNA colocalization specificity is driven by shared motifs and encoded protein function
- mRNAs with less compact folding, lower translation efficiency, and distinct protein-binding motifs are more likely to be in condensates
- Neurites have a particularly high degree of sequence and function-dependent mRNA organization

## INTRODUCTION

The spatial organization of biomolecules within cells is key to a wide variety of cellular functions, even though in many cases the determinants of this spatial organization remain poorly understood^1^. An important example is the compartmentalization of mRNAs into distinct subcellular regions, which can impact the function of the encoded protein by influencing the protein’s location, expression, and interactions^2–15^. However, we know little about the local protein and RNA environment of individual cytoplasmic mRNAs, or how these interactions impact mRNA regulation.

Biomolecular condensates, non-membrane-bound intracellular compartments enriched in specific proteins and nucleic acids, have emerged as a key aspect of RNA regulation^16^. Condensates can be in a relatively liquid physical state characterized by weak, transient interactions, or in a more gel or solid-like state, characterized by stronger, more stable interactions^17^. In many cases, RNA is necessary and sufficient for condensation, and RNA features impact condensate properties^18–23^. Recent evidence suggests that condensates can alter the physicochemical environment of their constituents, controlling internal hydrophobicity and pH^24–26^, melting nucleic acid duplexes^27^, and preferentially partitioning distinct RNA species^28^. Indeed, recent evidence indicates condensates may impact the function of mRNAs by selectively recruiting or displacing interacting molecules, altering localization, or changing the rate of enzymatic reactions^2,7–9,29–32^. Although condensation is a ubiquitous principle of intracellular organization, we are only beginning to understand how the condensate environment can impact gene expression.

Neurons require a high degree of spatial and temporal control over translation^33–35^, suggesting condensates could be well-suited to regulate neuronal mRNAs. A single neuron has 1,000 to 100,000 synapses that must each have a precise and continuously changing protein composition to make functional synaptic networks. One way neurons control their spatial proteome is by packaging mRNAs and associated RNA-binding proteins (RBPs) into condensates called transport granules, which move down neurites (dendrites and axons) for local translation when they reach the appropriate place and time^36–38^. Recent work indicates that 34 to 61% of synaptic spines are sites of active translation at any point in time^39^, and inhibiting local translation can impair the performance of mice in learning and memory tasks^13^. Moreover, transport granules are found in many other cell types, and cytoplasmic, mRNA-rich condensates with regulatory functions are still being actively discovered^3,40^. The condensate-regulated spatial distribution of mRNAs within cells thus appears to have vital functional consequences, especially in cells like neurons with complex spatial transcriptomes.

In contrast to their functional contributions in healthy neurons, condensates have also been implicated in neurodegenerative diseases. Pathological aggregates of RBPs (similar to those found in degenerating patient neurons) can be nucleated within condensates *in vitro* and in cultured cells, and this process can be exacerbated by disease-associated mutations^22,41–45^. Moreover, several neurodegeneration-associated genes, including TDP-43, tau, FUS, ATXN2, ANXA11, KIF5A, and hnRNPA2B1, play important roles in mRNA transport and local translation that can be impaired by disease-associated mutations^9,46–57^. Understanding the composition of RNA condensates, the interactions holding them together, and their roles in core gene-regulatory processes is thus not only important for elucidating basic cellular functions, but may also provide pivotal insights into neurodegenerative diseases.

One of the major challenges to understanding mRNA regulation and the role of condensates is the lack of technologies to map condensate-scale RNA organization genome-wide across subcellular regions. Insights have been gained through targeted approaches, such as APEXseq^58^, CLIP^59^, and FAPS^60^, that characterize RNAs associated with one protein of interest at a time. Prior untargeted methods that ligate RNAs together to make a chimeric molecule (e.g. PARIS, KARR-seq, RIC-seq) have elucidated RNAs that are bound together, revealing physical RNA-RNA interactions within tens of angstroms^61–65^. However, these approaches fail to capture emergent mesoscale (∼micron length) organization, as found in condensates. Insights into RNA organization can be gained through imaging spatial transcriptomics methods (e.g. MERFISH, seqFISH+), which give subcellular localization of single RNA molecules^66–68^. However, these methods require *a priori* knowledge of specific RNA targets and are constrained by inherent spatial resolution limits of confocal imaging, a challenge for characterizing RNAs densely clustered within condensates^69^. SPRITE (Split Pool Recognition of Interactions by Tag Extension) is a powerful tool to capture nucleic acid colocalizations on the scale of condensates (∼0.2 to 1 μm diameter)^70^. SPRITE has been validated to map RNA-DNA colocalization, identify colocalizations among abundant non-coding RNAs (ncRNAs) within known condensates, and discriminate between cytoplasmic and nuclear localized RNAs^71,72^. However, no previous methods, including SPRITE-based approaches, have interrogated genome-wide mature mRNA-mRNA spatial proximity.

Here we present RNA-SPRITE, an unbiased, sequencing-based method and computational approach for quantifying RNA-RNA colocalization, genome-wide across subcellular regions. We adapted previous SPRITE protocols^71^ to improve the reaction efficiencies and yield for less abundant mRNAs, enabling us to investigate drivers of specific mRNA colocalization and features associated with mRNA condensation. To gain insight into the strength and type of interactions controlling RNA condensation, we combined RNA-SPRITE with perturbations to disrupt or enhance condensation. Our results demonstrate genome-wide post-transcriptional RNA organization is ubiquitous and conserved between HEK cells and rodent neurons, with specific RNA features that are predictive of spatial colocalization. RNA-SPRITE provides unprecedented insights into global RNA organization, uncovering the mechanisms that shape it and its functional impact.

## RESULTS

### RNA-SPRITE detects colocalization among RNAs in both condensates and membrane-bound organelles

To interrogate RNA colocalization genome-wide across subcellular regions, we adapted prior protocols for SPRITE^73^ to develop RNA-SPRITE, optimizing for crosslinking and sonication with neurons/neurites and greater dynamic range of detection for lower abundance RNAs, through improved reaction efficiencies and increased yield (Methods). Briefly, we crosslink proteins and nucleic acids in live, intact cells using paraformaldehyde and disuccinimidyl glutarate (Figure 1A). We then lyse cells and sonicate into crosslinked clusters of proteins and nucleic acids. We can image the resulting clusters to confirm condensates remain intact and clusters have a maximum diameter of a few microns (Figure 1B). We next label the RNAs with cluster-specific barcodes using a split-pool method: clusters are distributed into a 96 well plate, with each well containing a unique oligonucleotide (“tag”) that is ligated onto all RNAs in that well (Figure 1A). All clusters are then pooled into a single tube and distributed into a new 96 well plate for a second oligonucleotide ligation. Repeating this process for 4 total rounds of ligation, plus the PCR adapter, creates a 5 tag barcode. Two molecules will only share the same barcode if they moved through the process together and thus were likely in close proximity in the live cell, resulting in colocalization within the same crosslinked cluster. We use next generation sequencing to read out the cDNA sequence and barcodes and align to the transcriptome. We see similar mRNA abundance in RNA-SPRITE data compared to RNAseq (Spearman ⍴ = 0.94, Figure S1G), indicating RNA-SPRITE does not have a strong bias for particular transcripts.

**Figure 1.**
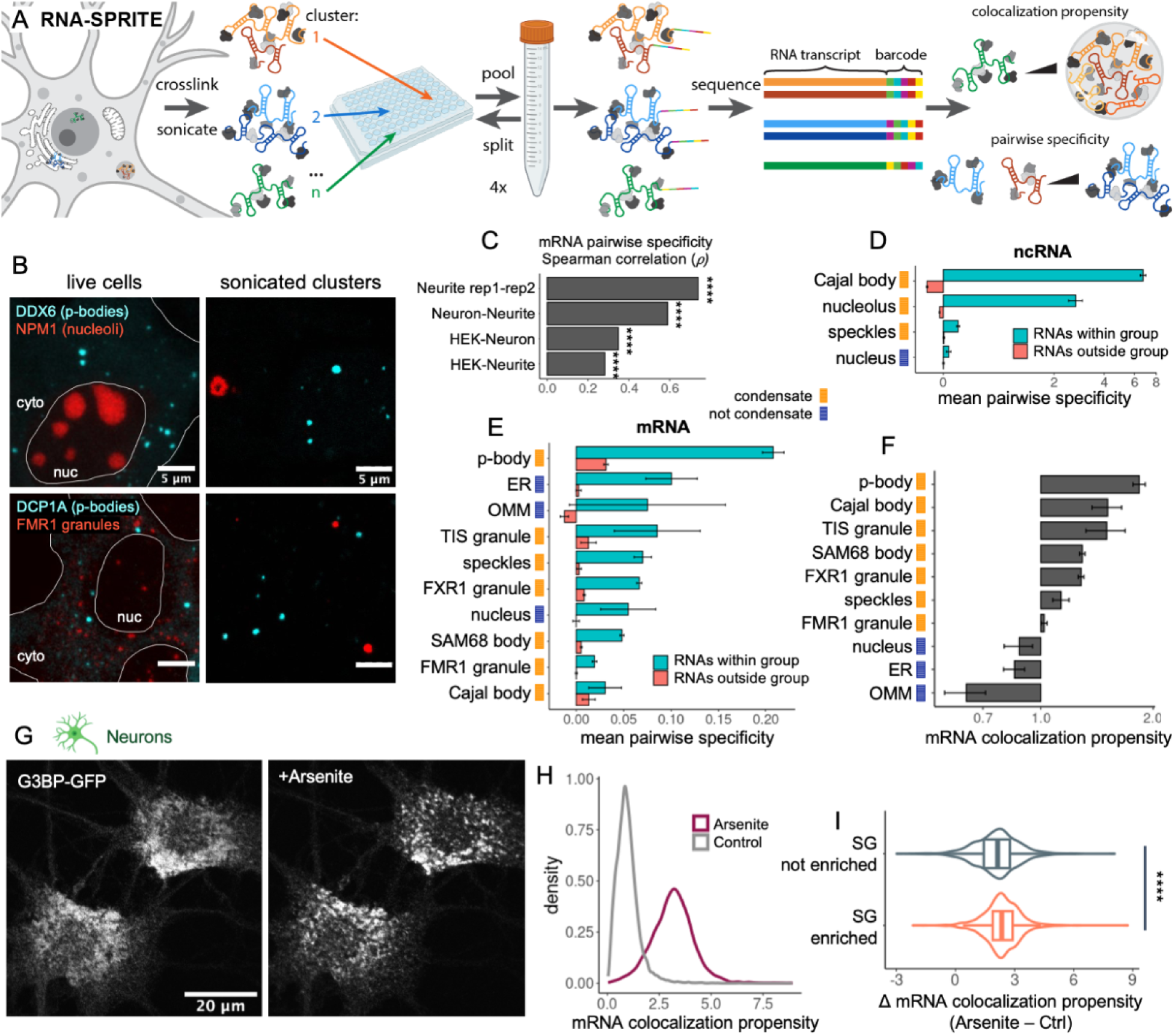
RNA-SPRITE detects colocalization among RNAs in both condensates and membrane-bound organelles. (A) RNA-SPRITE method schematic. (B) Live HEK cells and clusters after sonication labeled with DDX6-mEGFP, NPM1-mCherry, DCP1A-mEGFP, and FMR1-HaloJF585. (C) Spearman correlation of genome-wide, mature mRNA pairwise specificity. (D-E) For each ncRNA (D) or mRNA (E) in a given group (subcellular region), the mean pairwise specificity for all other RNAs in the same group or outside the group is plotted. (F) The mRNA colocalization propensity for each mRNA enriched in the indicated subcellular region is plotted. (G) Neurons were treated with sodium arsenite to induce stress granules, visualized with G3BP-GFP. (H) For all mRNAs, the relative probability density of mRNA colocalization propensity is plotted with and without sodium arsenite treatment. (I) The change in mRNA colocalization propensity after sodium arsenite is plotted for mRNAs annotated to be enriched in or depleted from stress granules (SG)^74^. Data presented as Tukey box plots with violin plots, Wilcoxon rank-sum test. (D-F) HEK cell data are presented as mean ± SEM, with arcsinh transform in (D) and log transform in (F) \*\*\*\**p* < 0.0001, See also Figure S1

To determine which pairs of RNA transcripts colocalized more than expected by chance, we developed a statistical framework capable of leveraging the large size of our datasets, and that is robust to noise and undersampling. Inspired by the widely-used single cell RNA-seq normalization method sctransform^75^, we used negative binomial regression (NBR) to model the number of colocalizations, which we define as the number of shared barcodes between each pair of RNAs. The NBR model predicts the expected number of colocalizations for a pair of RNAs, given the total colocalizations for each RNA and the genomic distance between the genes encoding the RNAs (to control for the co-transcriptional colocalization, Methods). Emulating sctransform, we calculate “pairwise specificity”: a measure of how much the observed number of colocalizations for a pair of RNAs deviates from that expected by chance, using the Pearson residual from the NBR model (Figure 1A and S1A-S1C).

To gain insight into RNA organization in both simple cells and cells with pronounced cytoplasmic heterogeneity, we performed RNA-SPRITE with human embryonic kidney 293T (HEK) cells, rat cultured primary cortical neurons, and isolated neurites from the cultured neurons (Figure S2D). The cultured neurons form a dense web of neurites, with extensive synapses, and fire action potentials spontaneously (Figure S2A-S2C), indicating that they are healthy and mature. We see a strong correlation of mRNA pairwise specificity values between replicates using independent NBR models (Spearman ⍴ = 0.74, Figure 1C). We also find good correlation among our independent HEK, whole neuron, and neurite datasets, with whole neuron and neurite datasets having the strongest correlation (Spearman ⍴ = 0.59), as expected. These results indicate that RNA-SPRITE is reproducible and robust, and that mesoscale mRNA organization is significantly conserved between HEK cells and rodent neurons.

We first examined whether RNA-SPRITE reliably detects known RNA localization patterns in both the nucleus and cytoplasm. We gathered data from studies using other methods, including APEXseq and FAPS, that individually characterized RNAs associated with distinct condensates and non-condensate regions^40,58,60,67,76–79^. Using the HEK RNA-SPRITE dataset, we found that RNAs annotated to be in the same subcellular structure indeed had more colocalizations with each other than with other RNAs: this holds for ncRNA (Figure 1D), but also much less abundant mRNAs (Figure 1E). For example, mRNAs annotated to be within p-bodies have, on average, nearly 7x higher mean pairwise specificity with all other p-body mRNAs than with non p-body mRNAs. mRNAs annotated as enriched in the ER also tend to colocalize more with other ER mRNAs, exhibiting ∼40x higher mean pairwise specificity. Based on these results, we conclude that RNA-SPRITE is able to accurately report RNA colocalization across a range of condensate and non-condensate subcellular regions in a single, untargeted assay.

Pairwise specificity reflects the likelihood that two RNAs exhibit frequent close spatial proximity to each other, which could occur in condensates and in non-condensate subcellular regions. We thus also sought a metric to quantify the propensity of each RNA to be concentrated within a region of high RNA density, as expected for RNA condensates. In our RNA-SPRITE data, we found some mRNAs colocalized much more with other mRNAs than would be expected based on their abundance. For each protein-coding gene, we quantified this mRNA colocalization propensity (i.e. apparent condensation) by dividing the number of observations shared with all other mRNA transcripts by the total number of observations for that mRNA, normalized so that the distribution mean is 1 (Figure 1A and S1C-S1D). Consistent with this metric reflecting high mRNA density, we found mRNAs annotated to be in known condensates had higher mRNA colocalization propensity, on average, than those annotated to be in non-condensate subcellular locations (Figure 1F).

To further examine whether our mRNA colocalization propensity metric reliably reflects condensation, we used sodium arsenite to cause mRNAs to condense into stress granules (Figure 1G). Upon arsenite treatment, we see an increase in mRNA colocalization propensity for nearly all mRNAs (Figure 1H), with mRNAs previously reported to be enriched in stress granules^74^ showing a greater increase in mRNA colocalization propensity than mRNAs reported to be depleted from stress granules (Figure 1I and S1E). The stress granule enriched mRNAs also had significantly greater pairwise specificity with each other after arsenite treatment than in untreated cells (Figure S1F). Adding to prior studies, these data suggest that most mRNAs are recruited to stress granules, but some are enriched more than others^74,80–82^. Together, our results indicate that mRNA colocalization propensity is a good metric of the tendency of an mRNA to be recruited to a condensate, and provides strong validation of RNA-SPRITE for quantifying both steady-state and perturbed RNA colocalization in living cells.

### RNA-SPRITE with hypotonic treatment probes reversibility of RNA colocalization genome-wide

Given the power of RNA-SPRITE for examining specific mRNA colocalization and condensation propensity, we deployed it to investigate the stability of interactions responsible for mRNA organization, using a simple hypotonic treatment approach. Upon addition of pure water to cell culture media, the osmotic pressure differential across the cell membrane drives water into the cells, increasing cell volume and diluting the cytoplasm (Figure 2A). Hypotonic treatment also causes ions and other small solutes to leave the cell, thereby decreasing electrostatic screening and disrupting weak interactions. These effects of hypotonic treatment are expected to partially destabilize liquid-like condensates^83–85^, since phase separation is directly dependent on the concentration of constituents and the strength of interactions.

**Figure 2.**
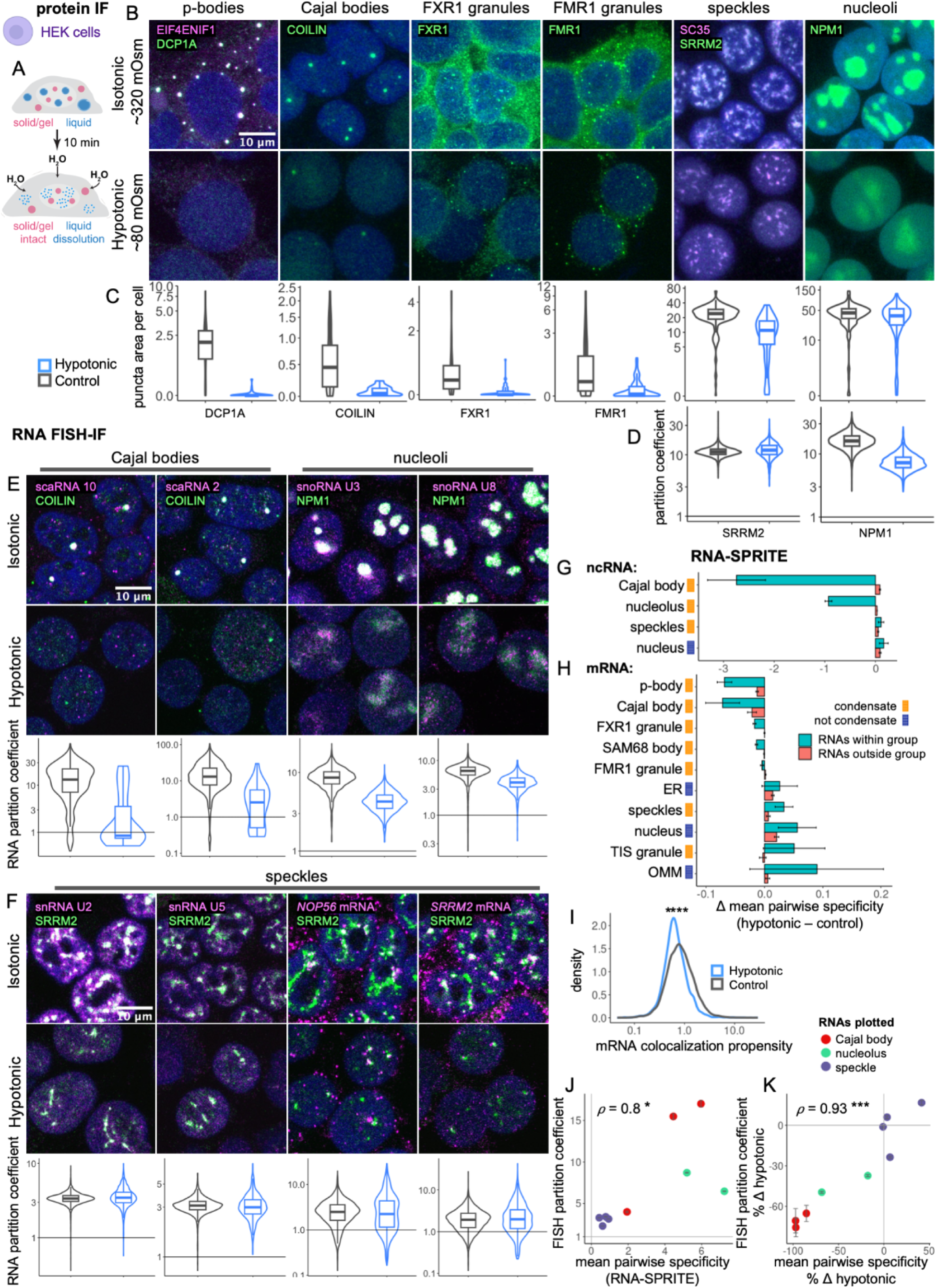
RNA-SPRITE with hypotonic treatment probes reversibility of RNA colocalization genome-wide. (A) Hypotonic treatment schematic. (B-D) IF z-stack maximum intensity projections (B) of condensates with and without hypotonic treatment with quantification of total condensate area per cell (C and Figure S3A) and partition coefficient of indicated marker into condensate (D). (E-F) FISH-IF images of condensate markers and enriched RNAs with and without hypotonic treatment with quantification of RNA partitioning into the condensate (separated channels Figure S3B). (G-H) For each ncRNA (G) or mRNA (H) in a given subcellular region, the mean pairwise specificity for all other RNAs in the same group or outside the group is plotted for the hypotonic treatment condition minus control, presented as mean ± SEM. (I) For all mRNAs, the relative probability density of mRNA colocalization propensity is plotted with and without hypotonic treatment; Wilcoxon rank-sum test reported. (J-K) Individual RNA partitioning into indicated condensate by FISH vs. RNA-SPRITE mean pairwise specificity with other RNAs in the same condensate in untreated conditions (J) and percent change in these values with hypotonic treatment (K). RNAs plotted are snoRNAs (U3, U8), snRNAs (U2, U4, U5, U6), and scaRNAs (2, 9, 10). Spearman correlation ⍴ reported. FISH data presented as mean ± SEM. (B, E-F) Nuclei stained with DRAQ5 or Hoechst, colored dark blue. (C-F) Data presented as Tukey box plots with violin plots. Arcsinh transform in (C) and log transform in (D-F). \**p* < 0.05, \*\*\**p* < 0.001, \*\*\*\**p* < 0.0001 See also Figure S3

We first validated this approach using immunofluorescence (IF) imaging. As expected, when we added three volumes of water to the media of live HEK cells for 10 min, we observed strong dissolution of p-bodies and Cajal bodies (Figure 2B-2C and S3A), suggesting that these bodies are held together by relatively weak interactions. Under the same conditions, we see partial dissolution of FXR1 granules, FMR1 granules, and speckles (Figure 2B-2C and S3A), which indicates they are held together by relatively stronger, more stable interactions. Interestingly, we find the total area of nucleoli does not change with hypotonic treatment, but the partitioning of constituents, such as NPM1, decreases (Figure 2B-2D). The scaffolding network of biomolecular interactions within nucleoli thus appears to remain largely in place, but the relative affinity of NPM1 for these nucleolar components decreases. Conversely, the total area of speckles decreases, but the partition coefficients of SRRM2 and SC35 do not change, suggesting that the affinity of these proteins for speckle components has not changed (Figure 2B-2D). These results demonstrate that differential effects of hypotonic treatment can report the reversibility of interactions within condensates.

To further examine the effects of this approach using hypotonic condensation disruption, we used Fluorescence In Situ Hybridization (FISH) in HEK cells to test if RNA condensation was affected by hypotonic treatment. In agreement with the protein imaging data, we found partitioning of small Cajal body-specific RNAs (scaRNAs) into Cajal bodies and small nucleolar RNAs (snoRNAs) into nucleoli was substantially reduced with hypotonic treatment (Figure 2E and S3B). Conversely, partitioning of small nuclear RNAs (snRNAs) and intron-rich mRNAs into speckles was not affected by hypotonic treatment (Figure 2F and S3B). These data provide further evidence that many Cajal body and nucleoli constituents are held together by relatively weak interactions, while speckles appear to be held together by stronger, more stable interactions.

We next investigated whether our RNA-SPRITE approach could provide genome-wide information about the effects of this hypotonic condensate dissolution. We subjected HEK cells to 10 min of hypotonic treatment before performing RNA-SPRITE and quantified mean pairwise specificity for groups of RNAs annotated to be in the same subcellular regions. For ncRNAs, we see a striking decrease in pairwise specificity among scaRNAs and among snoRNAs, just as we observed by imaging (Figure 2G). In close agreement with the imaging assays, we also see a large decrease in “within group” pairwise specificity for p-body or Cajal body mRNAs, while mRNAs enriched in FMR1 granules or FXR1 granules show a much smaller change (Figure 2H). Similar to snRNAs, speckle-enriched mRNAs do not have reduced colocalization with other speckle RNAs (Figure 2G-2H). As expected, “within group” pairwise specificity is not decreased for mRNAs enriched in non-condensate compartments: ER, nucleus, outer mitochondrial membrane (OMM). The non-condensate mRNAs appear to have greater pairwise specificity with each other because the number of colocalizations among them now deviates more from expectation based on the rest of the dataset, likely due to the loss of colocalizations among condensate mRNAs. These results provide further evidence that p-bodies and Cajal bodies are weakly bound by liquid-like interactions, while FMR1 granules and FXR1 granules have somewhat stronger gel-like interactions, and speckle components are even more strongly bound together.

To examine how hypotonic treatment affects mRNA condensation overall, we calculated mRNA colocalization propensity. As expected, we find a significant decrease (23% overall) in mRNA colocalization propensity after hypotonic treatment (Figure 2I). These results agree with the imaging data, and suggest that a specific subset of mRNAs enriched in liquid condensates become more diffuse upon hypotonic dilution, while many other mRNAs are not affected. Our mRNA colocalization propensity metric does not distinguish between condensates in liquid, gel, or solid states. However, mRNAs likely residing in liquid condensates can be putatively identified by their high sensitivity to hypotonic treatment (Figure S3C). Such transcripts are maintained in close spatial proximity through weak and readily reversible interactions. We investigated these mRNA transcripts to better understand what drives recruitment to liquid condensates, and we found significant enrichment of specific RBPs motifs (e.g. PUM1/2, ELAVL2/3, FUBP3, TIA1) in these sequences compared to hypotonic insensitive controls (Figure S3D). This indicates specific RBP motifs are involved in the recruitment of mRNAs to liquid condensates, in agreement with prior studies of individual RBPs^3,11^.

When we directly compare the data for individual RNAs, we see a significant correlation between RNA-FISH partition coefficient and “within group” RNA-SPRITE colocalization pairwise specificity in untreated conditions (Figure 2J). The partition coefficient reflects relative affinity, measured as the ratio of concentration (mean fluorescence intensity) of a molecule inside a condensate to outside. The percent change in partition coefficient with hypotonic treatment is highly correlated with percent change in RNA-SPRITE mean pairwise specificity (Spearman ⍴ = 0.93, Figure 2K). This suggests that similar to the partition coefficient, RNA-SPRITE colocalization pairwise specificity is a meaningful measure of relative molecular affinity. RNA-SPRITE allows us to measure molecular affinity changes with hypotonic treatment, and therefore, enables high-throughput quantification of the strength of forces holding RNAs together in condensates.

### mRNA colocalization hubs are revealed by RNA-SPRITE

Given the above validation of RNA-SPRITE for probing mesoscale RNA colocalization, we next sought to use it to generate a global map of cytoplasmic mRNA organization. To focus on mature cytoplasmic mRNAs and minimize confounds resulting from nascent nuclear mRNAs, we excluded barcodes that also mapped to known nuclear RNAs. We grouped the mRNAs into hubs based on colocalization pairwise specificity using singular value decomposition and Louvain community detection (Methods)^86^. Independent of hub assignment, we performed Uniform Manifold Approximation and Projection (UMAP)^87^ to visualize the hubs (Figure 3A).

**Figure 3.**
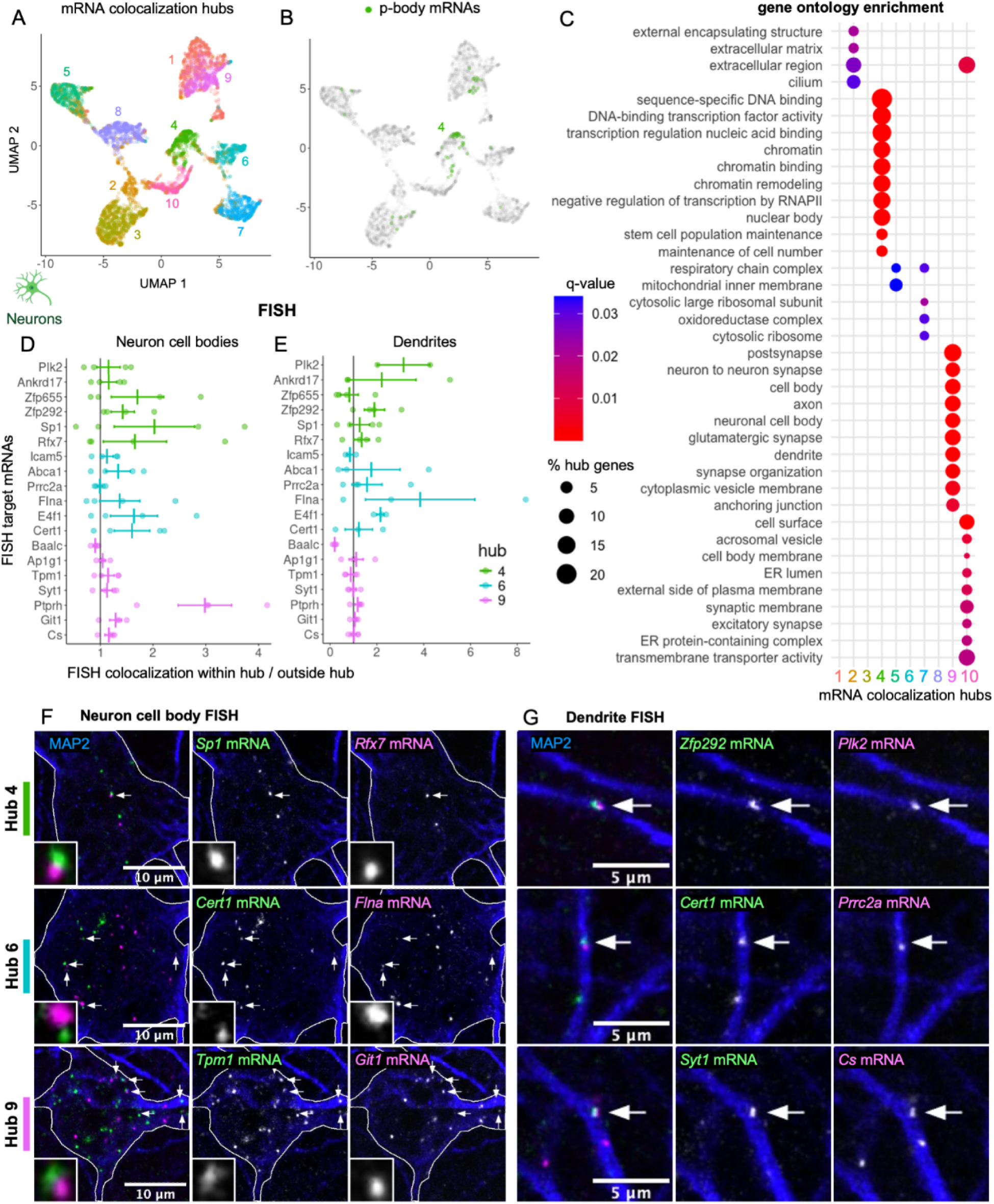
mRNA colocalization hubs are revealed by RNA-SPRITE. (A-B) Using whole neuron RNA-SPRITE data, mRNAs are plotted by UMAP coordinates and colored by colocalization hub (A) or enrichment in p-bodies (B). (C) Up to 10 enriched GO terms (q-value < 0.05) in each mRNA colocalization hub are plotted, where color reflects q-value and dot size reflects the percent of hub genes associated with each term. (D-E) Normalized pairwise colocalization of each mRNA with mRNAs in the same hub divided by median value outside of the hub in neuron cell bodies (D) or dendrites (E). Individual values and mean ± SE are shown. (F-G) IF-FISH images of neuron cytoplasm below nucleus (F) and dendrites (G) with arrows pointing to colocalizing mRNA transcripts and zoomed inset in (F). See also Figure S4

To investigate what the mRNA colocalization hubs represent, we first examined mRNAs that are enriched in p-body condensates^60^, and found that the majority are in hub 4 (Figure 3B), consistent with the hubs reflecting subcellular regions. We also observe distinct mRNA features in the hubs, including average abundance, transcript length, and guanine and cytosine (GC) nucleotide content (Figure S4A), suggesting mRNAs are not randomly distributed into the hubs. To further explore the hubs, we performed gene ontology (GO) analysis to see if hubs were enriched with genes that encoded proteins with similar function or localization (Figure 3C). Indeed, we found hub 4 was enriched in mRNAs encoding transcription and chromatin regulators, consistent with mRNAs previously reported as enriched in p-bodies^8,60^, and hub 9 was enriched in mRNAs involved in neuron-specific functions. We found hub 10 was enriched in mRNAs encoding proteins that were membrane-bound or secreted, indicating hub 10 likely reflects ER-associated mRNAs. Other hubs were enriched in mRNAs encoding cilia, secreted, mitochondrial, or ribosomal proteins. These data demonstrate a high degree of mRNA organization by protein function and localization.

We next sought to directly validate the RNA-SPRITE derived hubs by imaging RNA colocalization using FISH. We picked mRNAs from 3 hubs and quantified their relative colocalization frequency with each other, normalizing for abundance, with values greater than 1 indicating within-hub specificity (Methods). We found that most FISH targets from hubs 4 and 6 colocalized more with mRNAs from their own hub than those outside of their hub in both neuron cell bodies and in dendrites, while mRNAs in hub 9 only had higher within hub colocalization specificity on average in neuron cell bodies (Figure 3D-3G). However, we did not find all mRNAs in the same hub to colocalize more than expected by chance, suggesting that our clustering method is likely combining distinct subcellular compartments into a single hub.

### mRNAs with less compact folding and lower translation efficiency are more likely to be in mRNA condensates

The molecular features driving some mRNAs to be diffuse in the cytoplasm, while other mRNAs are enriched in endogenous cytoplasmic condensates (e.g., p-bodies, transport granules) are largely unknown. mRNA colocalization propensity quantifies the tendency of an mRNA to be enriched in regions with a high local mRNA concentration. Given that our calculated mRNA colocalization propensity is greater for mRNAs enriched in known condensates (Figure 1F) and is increased or decreased when we broadly enhance or disrupt condensation, respectively (Figure 1H and 2I), we interpret this metric as a readout of the tendency of an RNA to be in RNA-rich condensates, which could be in any physical state (i.e. liquid, gel, solid). We thus sought to use the metric to investigate the primary drivers of mRNA condensation across all cytoplasmic condensates.

We first considered how RNA folding may affect condensation by examining the percent of GC nucleotides in the sequence and length-normalized free energy of folding; high GC content is associated with more compact, stably folded RNA, which implies lower (more negative) free energy of folding. We found that percent GC content had a significant negative correlation with mRNA colocalization propensity across all three datasets (Spearman ⍴ = −0.26 to −0.3, Figure 4A-4B). Similarly, we estimated free energy of folding for each transcript using Nupack^88^, and again found that weaker intramolecular folding (less negative free energy of folding per kb) had a clear correlation with mRNA colocalization propensity (Figure 4A-4B). These data suggest that mRNAs with less compact folding are more likely to be in condensates, consistent with prior studies of individual mRNA species^16,89^, and the general property of extended, multivalent biomolecules to drive condensation^90–92^.

**Figure 4.**
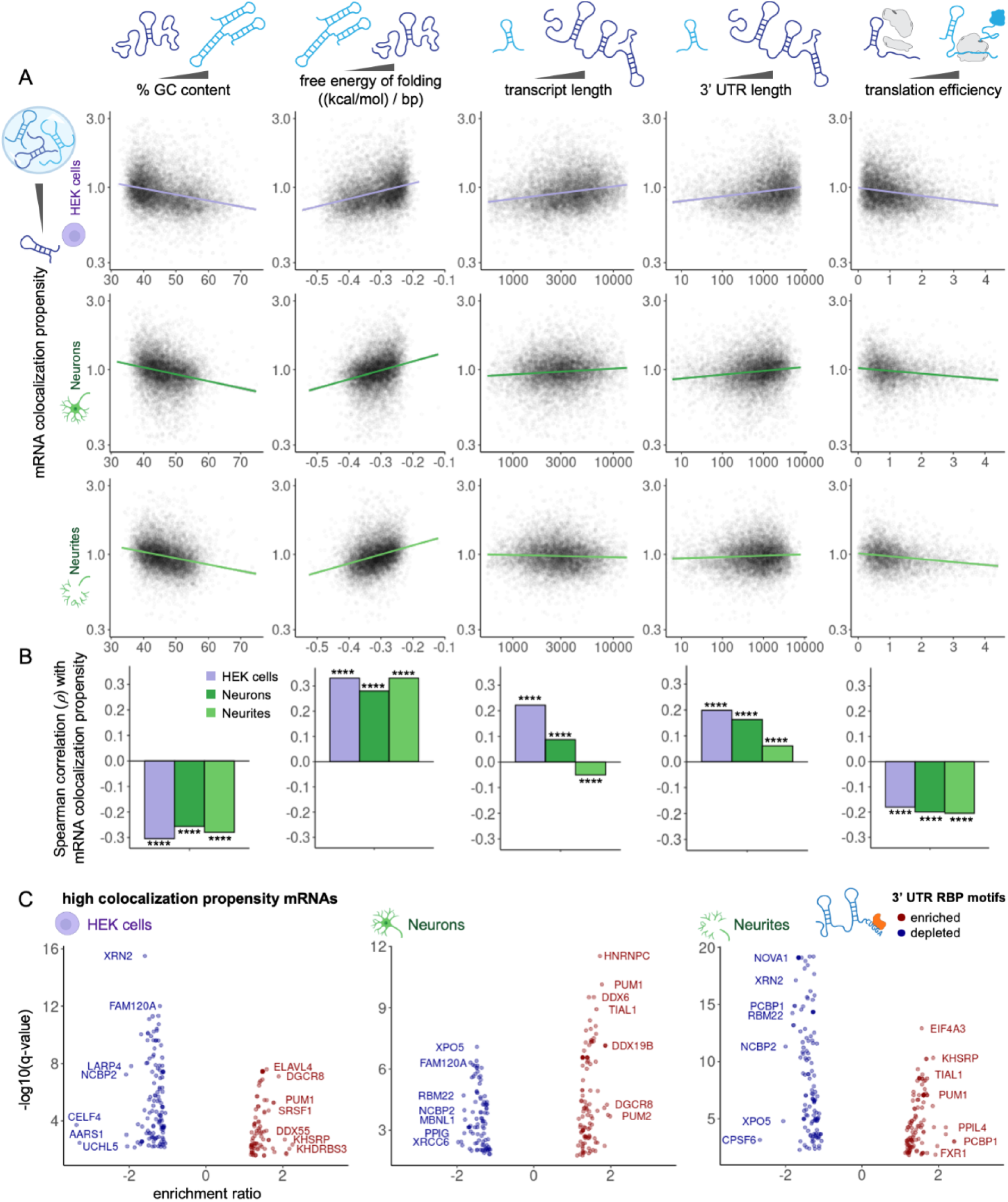
mRNAs with less compact folding and lower translation efficiency are more likely to be in mRNA condensates. (A) Individual mRNAs are plotted to compare the indicated metrics with mRNA colocalization propensity. Linear fits ± SE are plotted. Outliers removed; at least 98% of points shown. Colocalization propensity and lengths are log transformed. (B) Spearman correlation ⍴ plotted \*\*\*\**p* < 0.0001, no outliers removed. (C) RBP motif enrichment and depletion from the top quartile of high colocalization propensity mRNA 3’ UTRs vs. length-matched controls. Y-axis represents q-values < 0.05. See also Figure S5

mRNA regulation often occurs through interactions between RBPs and specific motifs in the 3’ untranslated region (UTR), whose accessibility could be impacted by RNA folding^93–96^. We thus sought to gain further insight into the relationship between RNA folding and condensation, and found free energy of folding per kb for coding sequences (CDS) and 3’ UTRs were both correlated with mRNA colocalization propensity (Figure S5A), again implying less compact mRNAs are more likely to be condensate-associated. However, CDS free energy of folding is more strongly correlated with mRNA colocalization propensity than 3’ UTR folding, across all 3 independent datasets, suggesting that 3’ UTR accessibility to RBPs is not the exclusive driver of mRNA condensation, in agreement with prior work^21^.

We next examined the importance of transcript length. Previous studies have shown that mRNA transcript length is correlated with stress granule enrichment^74,80,81^, however, it’s not clear if longer mRNAs are more enriched in endogenous condensates, in unstressed conditions. With RNA-SPRITE data we see a positive correlation between transcript length and mRNA colocalization propensity in HEK cells (Figure 4A-4B). However, this effect is much weaker in whole neurons and not observed in the neurite dataset. Examining mRNA subregions, 3’ UTR length has a more positive correlation with mRNA colocalization propensity compared to CDS length in all 3 datasets (Figure S5B).

These results suggest that RBP motifs in 3’ UTRs could play a role in recruiting mRNAs to condensates. We first confirmed from our sequencing data that all analyzed RBPs were expressed with greater than 5 transcripts per million (TPM) of their encoding mRNA (Figure S5D). We find that specific subsets of RBP motifs^97–100^ are enriched in the 3’ UTR of top quartile of high condensate propensity transcripts, versus length-matched controls (Figure 4C). Some of the RBPs are known to be enriched in condensates (e.g. DDX6, FXR1, TIAL1, PUM1). Conversely, we also find specific RBP motifs that are relatively depleted from the top quartile high mRNA colocalization propensity transcripts (Figure 4C). Future work will be needed to fully elucidate the relationships between RBP interactions and mRNA condensation, however, these data demonstrate that specific RBP motif interactions likely influence the recruitment of mRNAs to condensates or maintain their diffuse distribution in the cytoplasm.

The poorly understood relationship between condensation and translation is of intense interest in the field^101^. To investigate this question genome-wide, we collated translation efficiency data derived from dividing ribosome-associated sequencing reads by total reads in HEK cells^102^, neuron cell bodies^103^, and neurites^103^. We also considered translation initiation efficiency data^104^ in HEK cells (Figure S5C). We found a consistent, significant correlation in all 3 cellular contexts, where transcripts with higher mRNA colocalization propensity tend to have lower translation efficiency (Spearman ⍴ = −0.20 to −0.23, Figure 4A-4B). Our data suggest that, overall, mRNAs in condensates have lower translation efficiency than those diffuse in the cytoplasm.

### mRNA colocalization pairwise specificity is driven by shared RBP partners and encoded protein interactions

In the previous section we investigated global drivers of mRNA condensation. We can also use RNA-SPRITE to elucidate universal factors that promote two mRNAs to specifically colocalize with each other inside or outside of condensates. We examined three possible drivers of mature, cytoplasmic mRNA colocalization pairwise specificity: RNA-RNA duplexes, encoded protein interactions, and shared RBP motifs. mRNA localization may be influenced by specific physical contacts between separate mRNA molecules, stabilized by Watson-Crick base pairing and/or non-Watson-Crick interactions^105^, which we will refer to collectively as “duplexes”^63^. Alternatively, nascent proteins still being translated from their mRNAs can form stable interactions and thus alter mRNA localization^106^. Finally, RBPs exert enormous control over mRNAs, and have been suggested as the primary mediators of mRNA localization. The degree to which these factors direct cell-wide mRNA organization is not well understood.

We first examined the impact of direct intermolecular mRNA-mRNA duplexes on genome-wide mRNA colocalization, by incorporating data from proximity ligation studies^63,64,107,108^. Using our NBR framework, we can add additional terms to the baseline model for each dataset, and estimate corresponding coefficients to quantify how well different features predict mRNA colocalization (Methods). The model parameter α reflects a multiplicative change in mRNA-mRNA colocalizations (α > 1 implies more colocalizations, while α < 1 implies fewer). Mature mRNA pairs from different genes annotated to physically interact had 30% more colocalizations on average (“mRNA duplex multiplier,” α = 1.30) in HEK cells, but only 4% more in neurites (α = 1.04) and were not more likely to colocalize in whole neurons (α = 1.00; Figure 5A). These differences may result from the fact that proximity ligation data is from human and mouse cell lines, while our neuron data is from rat primary cells. These results suggest specific duplexes between mature mRNAs can contribute to RNA colocalization in human cell lines, however, they are not broadly conserved.

**Figure 5.**
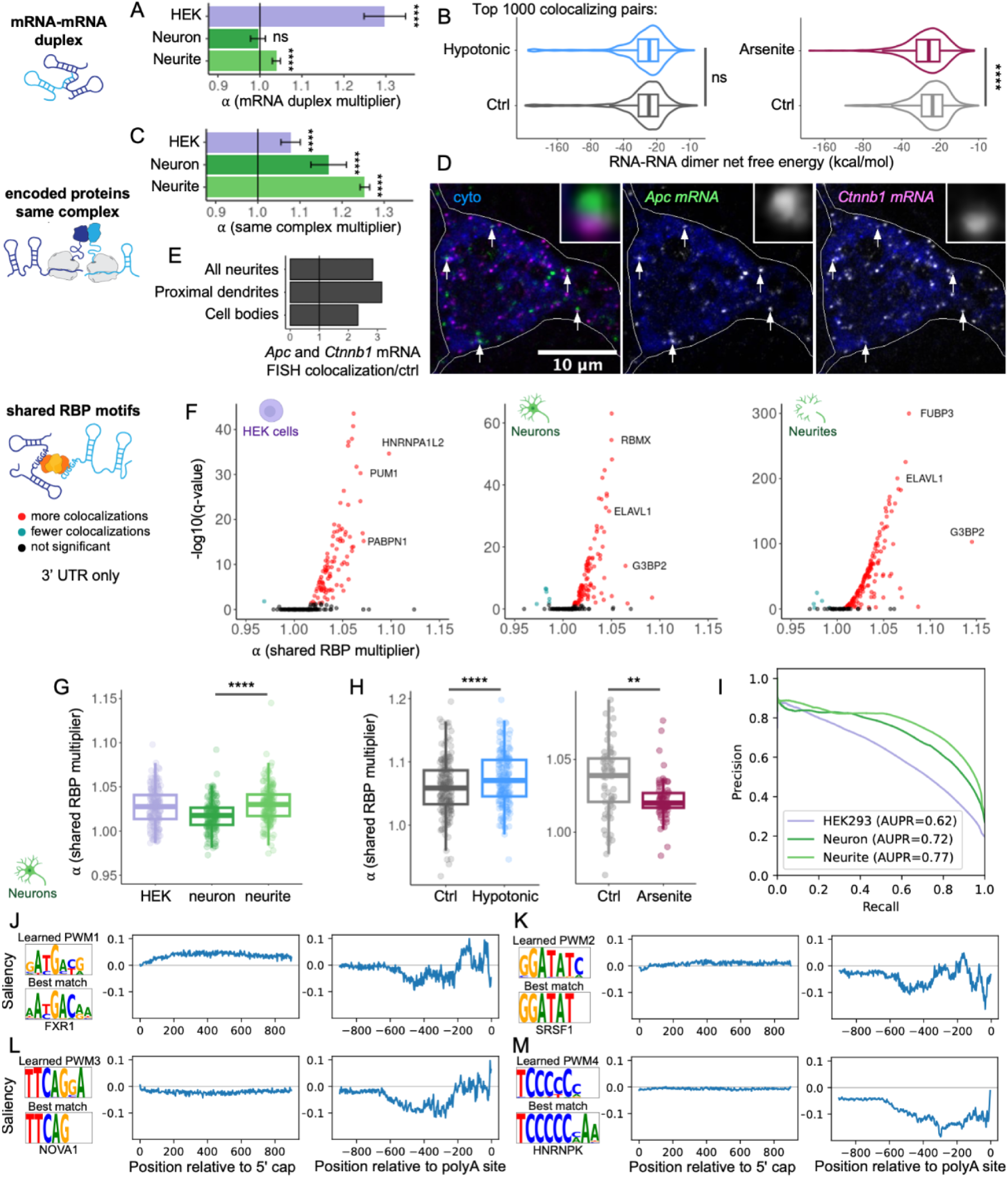
mRNA colocalization pairwise specificity is driven by shared RBP partners and encoded protein interactions. (A) The multiplicative effect on mRNA pairwise specificity (α) if two mRNAs are annotated to form a duplex^107^. (B) Free energy of folding for intermolecular dimers minus values for separate monomers. Highest confidence 1000 colocalizing pairs in treated datasets vs. those in untreated, presented as Tukey box plots with violin plots and arcsinh transform. (C) The multiplicative effect on mRNA pairwise specificity (α) if two mRNAs encode proteins that physically interact. (D) Two mRNAs encoding proteins in the same complex imaged via IF-FISH in a neuron, with UPF1 antibody (blue) marking the cytoplasm below the nucleus and zoomed insets. (E) Normalized FISH colocalization of *Apc* and *Ctnnb1* mRNAs over normalized colocalization with control mRNAs. (F) For each RBP, the multiplicative effect on mRNA pairwise specificity (α) if two mRNA 3’ UTRs share a motif for the RBP vs. Benjamini-Hochberg adjusted significance is plotted. (G-H) The RBP multiplicative effect on mRNA pairwise specificity (α) for each dataset (G) and with perturbations (H). RBPs with a significant, positive effect in at least one dataset are plotted along with Tukey box plots. (I) RNA sequence-based deep learning model performance, Area Under Precision-Recall (AUPR) (J-M) Position weight matrices (PWMs) predicting neuron RNA pairwise specificity with transcript location effect and best match known RBP motif. (B, G-H) Wilcoxon rank-sum test (A and C) Error bars represent 95% confidence intervals \*\**p* < 0.01, \*\*\*\**p* < 0.0001, ns = not significant See also Figure S6

To further investigate whether mRNA-mRNA duplexes drive specific colocalization, we used Nupack^88^ to estimate the free energy of heterodimerization of RNA pairs. As a positive control, we observed colocalizing mRNA pairs found to form a duplex by proximity ligation experiments had significantly lower free energy of dimerization versus non-colocalizing length-matched controls (Figure S6A), as expected. However, we did not observe a difference in free energy of dimerization between colocalizing and non-colocalizing pairs in our 3 untreated datasets (Figure S6B) or between the top colocalizing pairs with hypotonic treatment versus control (Figure 5B and S6C). Therefore, specific mRNA-mRNA duplexes likely are not a prevalent driver of mature mRNA colocalization under typical conditions. Interestingly, however, we do see significantly more stable predicted mRNA-mRNA duplexes (lower free energy of dimerization estimates) with arsenite treatment versus control (Figure 5B and S6D), suggesting that mRNA-mRNA duplexes are substantially increased during stress granule condensation, in agreement with prior work^23,109,110^.

We next examined whether mRNA-mRNA colocalization is significantly promoted by interactions of their encoded proteins. Again using our NBR framework, we found that two mRNAs encoding proteins annotated to be in the same protein complex^111–113^ have 8% more colocalizations on average (“same complex multiplier,” α = 1.08) in HEK cells, 17% more (α = 1.17) in whole neurons, and 25% more (α = 1.25) in neurites than otherwise expected (Figure 5C), but not with randomized negative control annotations (Figure S6E). We used FISH to validate colocalization of two mRNAs in neurons that encode interacting proteins: *Apc* and *Ctnnb1* (Figure 5D-5E). Our data thus suggest that it is common for mRNAs with shared function to colocalize, and this may be in part due to co-translational interactions.

Finally, we asked whether sharing an RBP binding partner increased the observed colocalizations between two mRNAs. For each of 209 expressed RBPs (Figure S5D), we identified putative binding sites in each transcript^97–99^ and fit an NBR model to estimate the percent increase in colocalizations between mRNA pairs that share the RBP partner (Methods). We found many RBPs predict increased colocalizations between their mRNA targets (“shared RBP multiplier,” α > 1), when we looked at motifs in 3’ UTRs (Figure 5F) or pairs of motifs within 200 nt of each other in the whole transcript (Figure S6G). For example, two mRNAs sharing a 3’ UTR motif for G3BP2, a protein constituent of neuronal transport granules involved in axon repair^114^, have 14% more colocalizations (α = 1.14) in neurites, on average, than expected. Interestingly, we see that sharing an RBP site predicts more colocalizations between a pair of mRNAs in neurites than it does in whole neurons, on average (Figure 5G). This likely results from neurite mRNAs typically being incorporated into transport granules. The frequency at which mRNAs are transported together versus individually has been a topic of debate within the field^115^, however, our data are more consistent with RBPs commonly recruiting multiple target mRNAs into a single transport granule^116^.

Sharing an RBP partner or encoding proteins that physically interact can increase colocalizations between two mRNAs, but the findings described above do not address whether these effects are significant drivers of mesoscale condensation. To address this question, we leveraged our experiments with hypotonic and arsenite treatment to cause diminished or enhanced mRNA condensation, respectively. These perturbations do not significantly affect colocalizations between mRNAs encoding proteins that physically interact (Figure S6F). Moreover, when we disrupt weak interactions with a hypotonic treatment, we find shared RBP motifs become even more predictive of mRNA pairwise specificity, indicating most RBP-motif interactions are relatively stable, even with widespread condensate dissolution (Figure 5H). Conversely, when we broadly increase mRNA condensation via arsenite-induced stress granules, sharing an RBP partner has a smaller effect on mRNA pairwise specificity (Figure 5H), indicating that other types of interactions become more dominant. Indeed, stress granule-enriched RBPs^117^ do not have greater mRNA colocalization predictive power (α) versus other RBPs following stress granule formation (Figure S6H). These data suggest that, overall, co-translational protein interactions and specific interactions of RBPs with distinct mRNA sequence motifs exist both inside and outside of condensates, but may not be dominant in driving liquid-like condensate formation.

To further explore the sequence determinants of mRNA colocalization, we developed a deep learning model to predict pairwise specificity from the sequences of the two mRNAs (Figure S6I, Methods). We found transcript sequence alone is highly predictive of mRNA pairwise specificity (test set AUPR = 0.62 to 0.77), with the model trained on the neurite dataset demonstrating the best precision-recall performance (Figure 5I). We identified informative sequence regions by calculating saliency, the gradient of model output with respect to the sequence input: positive saliency predicts higher pairwise specificity, while negative saliency predicts lower pairwise specificity^118–120^. The model was able to learn *de novo* sequence motifs that affected the likelihood of mRNA colocalization when shared by both transcripts, and also gave insight into the significance of motif localization (Figure 5J-5M). Our model uncovered motifs, many of which correspond to known RBP motifs, that predicted greater pairwise specificity if within 1kb of the 5’ cap (Figure 5J-5K) or within 200 nt of the polyA site (Figure 5J-5L). We also found motifs that predicted lower pairwise specificity within 1kb of the 5’ cap (Figure 5L) or within 200 nt of the polyA site (Figure 5M). These deep learning analyses demonstrate that subcellular mRNA organization has prominent sequence-encoded organizing principles, including individual RBP motifs.

## DISCUSSION

The local microenvironment of an mRNA can impact its expression, processing, localization, degradation and encoded protein function^101,121,122^. Recent findings suggest condensates are ubiquitous RNA-rich microenvironments throughout living cells. Here, we’ve developed and deployed RNA-SPRITE as a genome-wide map of RNA spatial proximity. We find evidence of extensive mesoscale mRNA organization (Figure 3) that is significantly conserved between HEK cells and rodent neurons (Figure 1C), with neurites having a particularly high degree of interpretable mRNA arrangement (Figure 5). Our results suggest that interactions that bring specific mRNAs together are distinct from those that promote condensation. The specificity of mRNA colocalization is strongly influenced by shared RBP motifs and encoded protein function, while mRNA condensation is associated with less compact folding, lower translation efficiency, and specific RBP partners (Figure 6).

**Figure 6.**
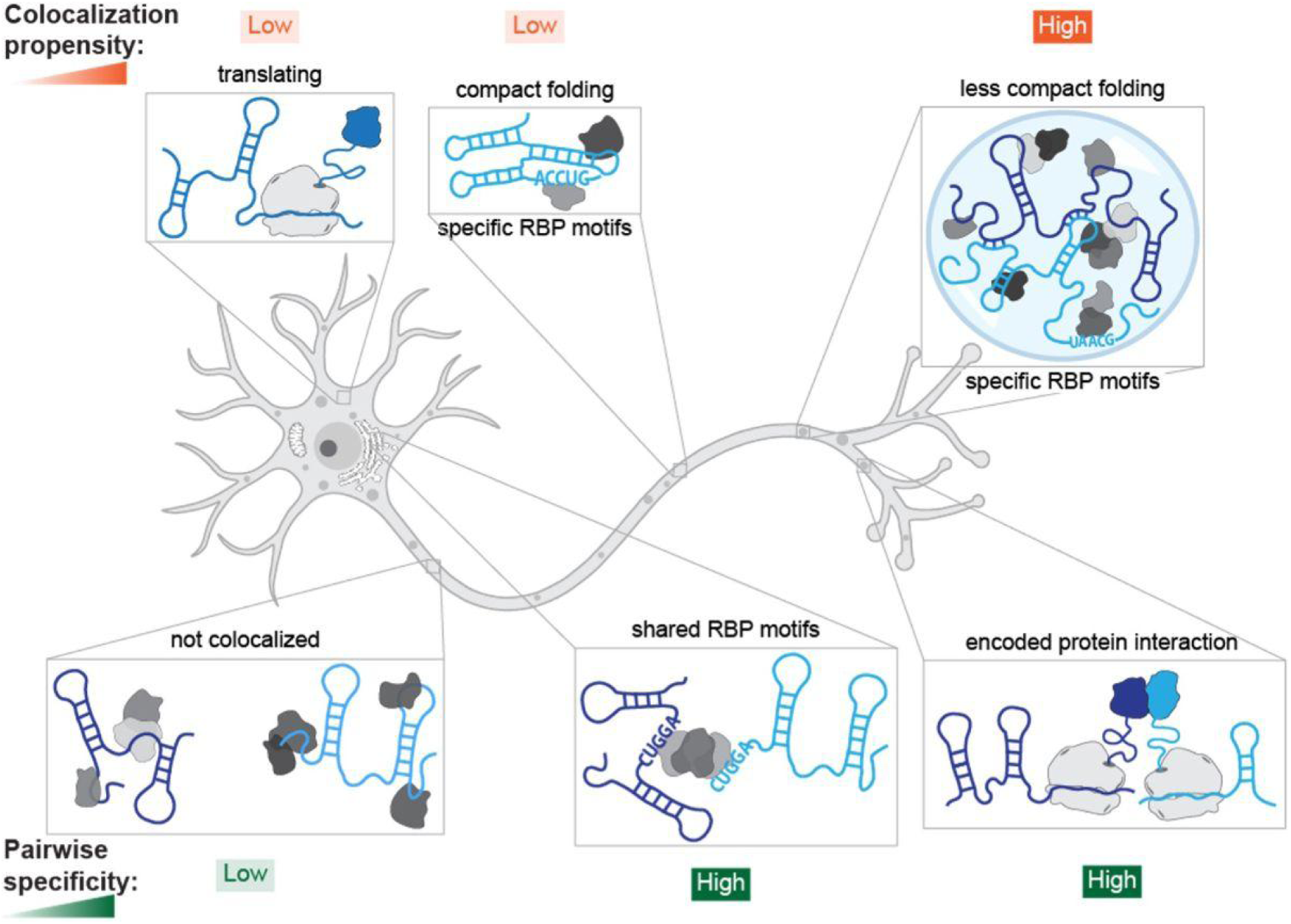
RNA-SPRITE reveals drivers of RNA condensation and colocalization specificity. mRNAs with high colocalization propensity (likely in condensates) tend to have lower translation efficiency and less compact folding. Distinct RBP motifs are associated with either low or high colocalization propensity mRNAs. mRNA colocalization pairwise specificity is driven by shared RBP motifs and shared function of encoded proteins. We observe all of these trends in HEK, whole neuron, and neurite datasets, with zoomed-in windows highlighting examples of potential subcellular locations.

Our condensate perturbation experiments provide further evidence that mRNA pairwise specificity and condensation propensity have distinct molecular drivers. Hypotonic treatment is less disruptive of RBP motif-driven or encoded protein-driven mRNA colocalization, even while it does notably dissolve liquid condensates, through disruption of a weaker set of interactions holding them together. This suggests these interactions remain largely intact outside of condensates, in agreement with prior work investigating several different *in vitro* reconstituted RBP-RNA condensates^123^. Conversely, increasing mRNA condensation through arsenite-induced stress granule formation is associated with increased RNA-RNA duplexes, while RBP-motif interactions play a smaller role. Specific RBP motifs can promote or inhibit recruitment to condensates, but broad condensate perturbations do not appear to affect motif-RBP interactions. Our results thus support a model where stable RBP motif interactions are not the dominant mechanism holding liquid condensates together. Instead, individual mRNAs and their stably-bound protein partners represent a foundational unit that can act as a building block for larger-scale, dynamic condensate assembly, through multivalent interactions among RNAs and proteins to maintain the cohesion of liquid condensates^121^.

RNAs can play essential structural roles in condensates^16,21,89,110,124,125^, however, the impact of intermolecular RNA interactions and RNA structure on condensation and subcellular organization are not well understood. We observe that experimentally annotated mature mRNA-mRNA duplexes reliably form in human cell lines, but are not well conserved in rodent neurons. Our free energy of dimerization estimates suggest mRNA-mRNA duplexes are not a dominant driver mRNA organization in untreated conditions, but do become more prevalent during stress granule induction, in agreement with prior studies^23,109,110^. Thus stable, specific heterodimer mRNA-mRNA duplexes can contribute to mRNA organization, but are likely not key contributors to the ubiquitous mesoscale mRNA organization we observe with RNA-SPRITE. When we examine colocalization propensity, we find mRNAs with less compact folding are more likely to be in condensates. This suggests the larger surface area and more accessible single-stranded segments promote multivalent interactions with other RNAs or proteins, in agreement with studies of individual mRNAs^89,126^. Our evidence of enhanced multivalency in condensate mRNAs is consistent with mRNAs routinely contributing to the structure of condensates, instead of simply being passive constituents^127^.

The relationship between mRNA condensation and translation is an outstanding question in the field^122^. Many condensates contain translationally repressed mRNAs, inhibiting translation increases mRNA condensation, and condensate dissolution is associated with increased translation^60,122,128,129^. Conversely, it has been shown that translation can occur, and may even be promoted, in some condensates^2,3,15,130–133^. In the mammalian cellular contexts we examined, overall, lower translation efficiency mRNAs were more likely to be in condensates. This result does not contradict findings demonstrating active translation in specific condensates, but does suggest translation is more likely to occur outside of condensates.

Interestingly, we find organization of mRNAs by the function of their encoded protein is widespread. We observe mRNAs encoding proteins that physically interact have substantially more colocalizations with each other (Figure 5C), especially in neurites, and mRNA colocalization hubs are enriched in specific GO terms (Figure 3C). Functionally related mRNAs could be brought together by shared RBP partners or by interactions of their encoded nascent polypeptides during translation. Co-translational interactions can influence mRNA localization^106^ play crucial roles in encoded protein function^134^, and even occur on translationally stalled transcripts^135^. Neurons have many transcripts inhibited during translation elongation that can be packaged together with translation machinery into transport granules^136–138^. Sorting of functionally related mRNAs into the same transport granules through co-translational interactions or shared RBP partners could facilitate coordinated transport and local translation in neurons.

In conclusion, RNA-SPRITE has enabled us to achieve unique insight into global subcellular mRNA organization and the factors that drive it. RNA-SPRITE provides a powerful tool for (1) uncovering novel condensates and specific mRNA recruitment to other subcellular regions, (2) delineating the biomolecular interactions that underlie RNA organization and condensation, and (3) providing new insights into the mechanisms of RNA regulation and dysregulation in disease. Building off of the findings presented here, future work leveraging RNA-SPRITE has the potential to discover modulators of RNA organization, including therapeutic approaches to pathologies such as neurodegeneration diseases, where the dysregulation of mRNA-rich condensates appears to play a central role.

### Limitations of the study

RNA-SPRITE is potentially constrained by undersampling, transcript discrimination, and quality of annotation data. Deeper datasets could help reduce colocalization false negatives and refine identification of colocalization hubs corresponding to distinct subcellular compartments. Also, we do not report colocalization between copies of the same transcript sequence or between transcripts with high sequence homology because it is difficult to determine if the two sequences came from the same RNA molecule or separate molecules. Improved experimental methods and algorithms to predict RBP binding, RNA folding, and mRNA-mRNA duplexes would enhance our insights into how these factors control mRNA colocalization and condensation. Such characteristics of individual RNAs are difficult to predict, however, when we look at these features genome-wide we can identify high confidence trends.

## ACKNOWLEDGMENTS

We thank Mitchell Guttman, Isabel Goronzy, and Mario Blanco for SPRITE oligonucleotides, protocols, and scripts; Lennard Wiesner and Lifei Jiang for oligonucleotides and FISH support; Shree Tanneti, Andrew Esteves, Kumar Mritunjay, Archana Sharma, Jean Schwarzbauer, and Lynn Enquist for sharing rodent embryos; Jennifer Miller, Wei Wang, and the Genomics Core Facility for genomics support; David Sanders and Lian Zhu for plasmids; Evangelos Gatzogiannis for microscopy assistance; Elizabeth Gavis, Amy Strom, Anita Donlic, and Nima Jaberi-Lashkari for thoughtful comments on the manuscript; Britt Adamson for useful advice on the figures and project narrative; and other members of the Brangwynne Lab for helpful feedback, discussions, and experimental support. This work was supported by the NIH NIDA (R21DA056345), the Howard Hughes Medical Institute (HHMI), the Princeton Biomolecular Condensate Program, the Princeton Center for Complex Materials, a MRSEC (NSF DMR2011750), the St. Jude Collaborative on Membraneless Organelles, and the AFOSR MURI (FA9550-20-1-0241), and the Chan Zuckerberg Initiative Exploratory Cell Network. L.A.B. was supported by an HHMI Helen Hay Whitney Fellowship. S.A.Q. is supported by an HHMI Hanna H. Gray Fellowship.

## AUTHOR CONTRIBUTIONS

L.A.B., S.A.Q., C.P.B., and D.A.K. designed the study. L.A.B. and S.A.Q. performed the experiments. L.A.B., D.A.K., S.A.Q., and T.J.C. developed data analysis methods and analyzed the data. O.K. provided advice and scripts for the free energy of folding analyses. D.A.K. advised the analysis. C.P.B. advised the experiments and analysis. L.A.B. and C.P.B. wrote the manuscript with input from all authors. L.A.B. made the figures with contributions from all authors.

## Data and code availability

The sequencing data generated in this study have been deposited in the Sequence Read Archive (NCBI: SRA) under accession code BioProject ID PRJNA1241813. Custom scripts are available at: https://github.com/davidaknowles/adaglm, https://github.com/davidaknowles/locnet, and https://github.com/SoftLivingMatter/RNA-SPRITE-Becker-2025.

## Supplemental figures

**Figure S1.**
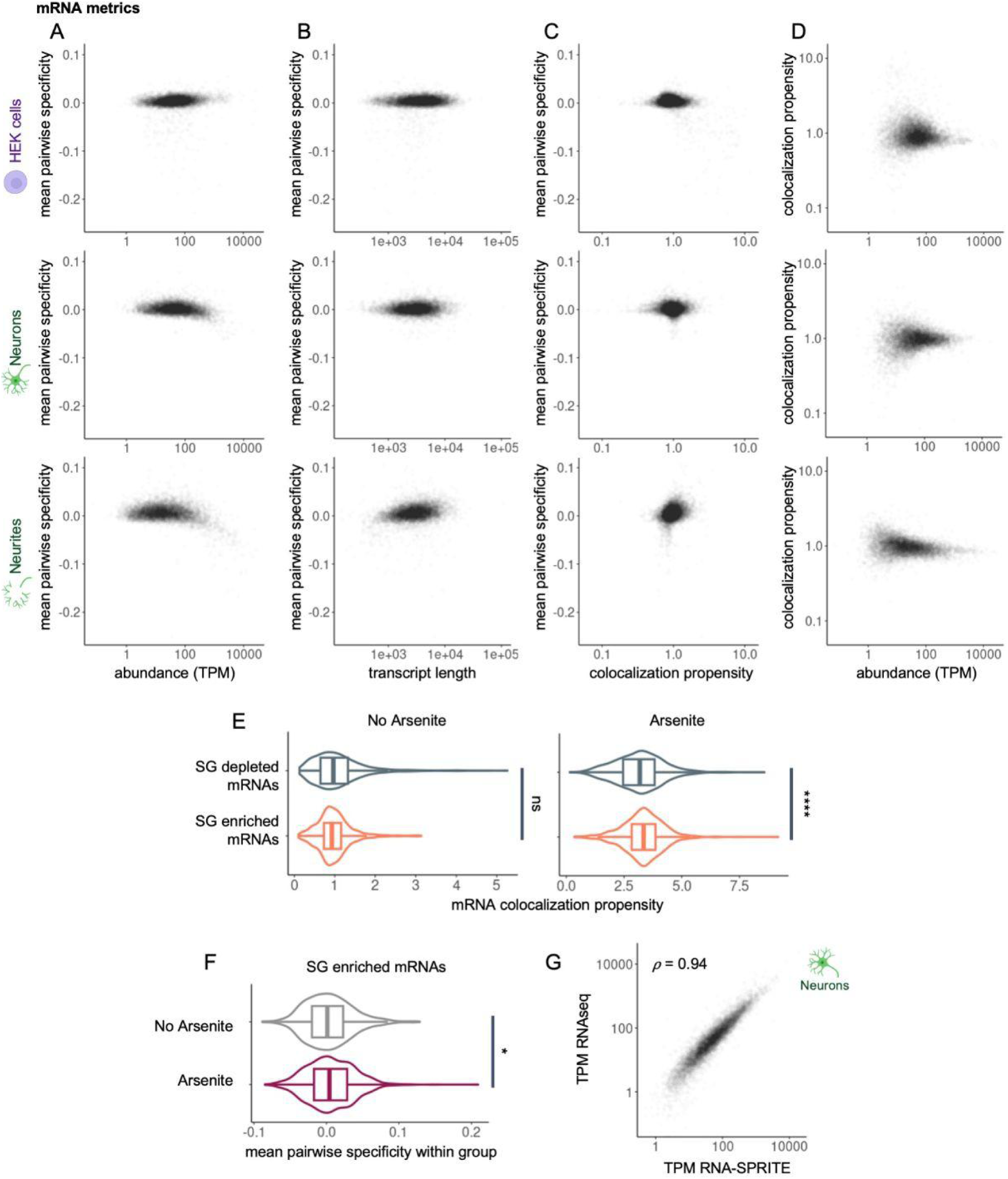
Characterization of RNA-SPRITE colocalization propensity and pairwise specificity metrics for mRNAs, related to. Figure 1 (A-D) Individual mRNAs are plotted to show correlations among transcripts per million (TPM), transcript length, mRNA colocalization propensity, and mean pairwise specificity for each mRNA with all other mRNAs. (E) The mRNA colocalization propensity for stress granule (SG) depleted and enriched mRNAs^74^ before and after arsenite treatment. (F) The mean pairwise specificity for SG enriched mRNAs with all other SG enriched mRNAs before and after arsenite treatment. (G) Individual mRNAs are plotted with TPM measurements from RNA-SPRITE vs. RNAseq in cultured neurons. (A-D and G) TPM, length, and colocalization propensity axes are log transformed.

**Figure S2.**
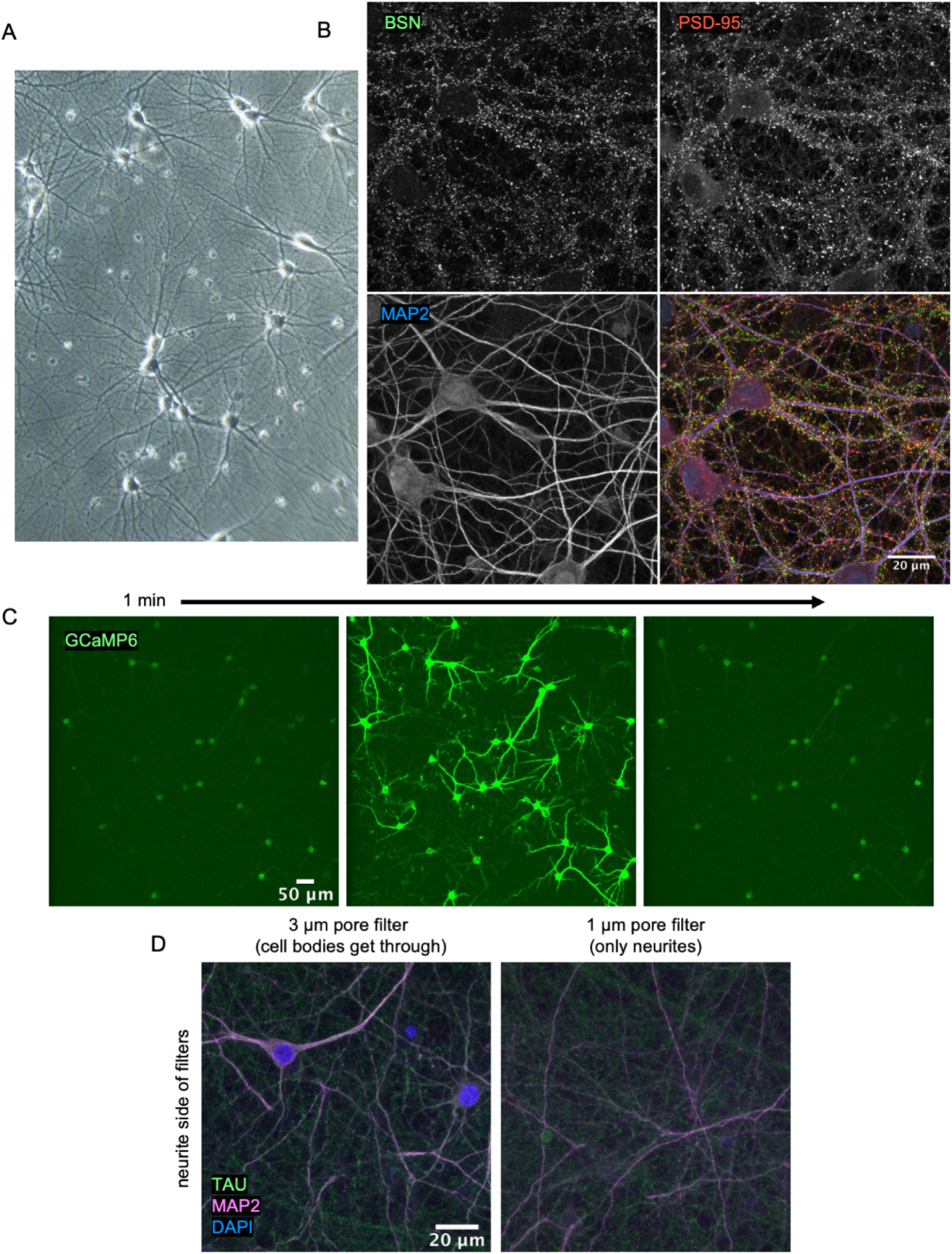
Rat primary neuron cultures form functional, mature, electrically active synapses. (A) Day 18 in vitro cultured cortical neurons have a dense web of neurites and are morphologically mature. (B) Extensive overlap of antibody stains for presynaptic marker BSN and postsynaptic marker PSD-95 indicates abundant functional synapses.(C) The genetically encoded calcium indicator GCaMP6 displays spontaneous spikes that are well synchronized among the neurons, demonstrating that the cells are electrically active and innervating each other. (D) Neurons were grown on hanging filters (Methods) to isolate neurites. Unlike the 3 µm pore filters, on the neurite only side of the 1 µm pore filters, we see evidence of axons and dendrites via TAU and MAP2 antibody labeling, but not cell bodies labeled with DAPI.

**Figure S3.**
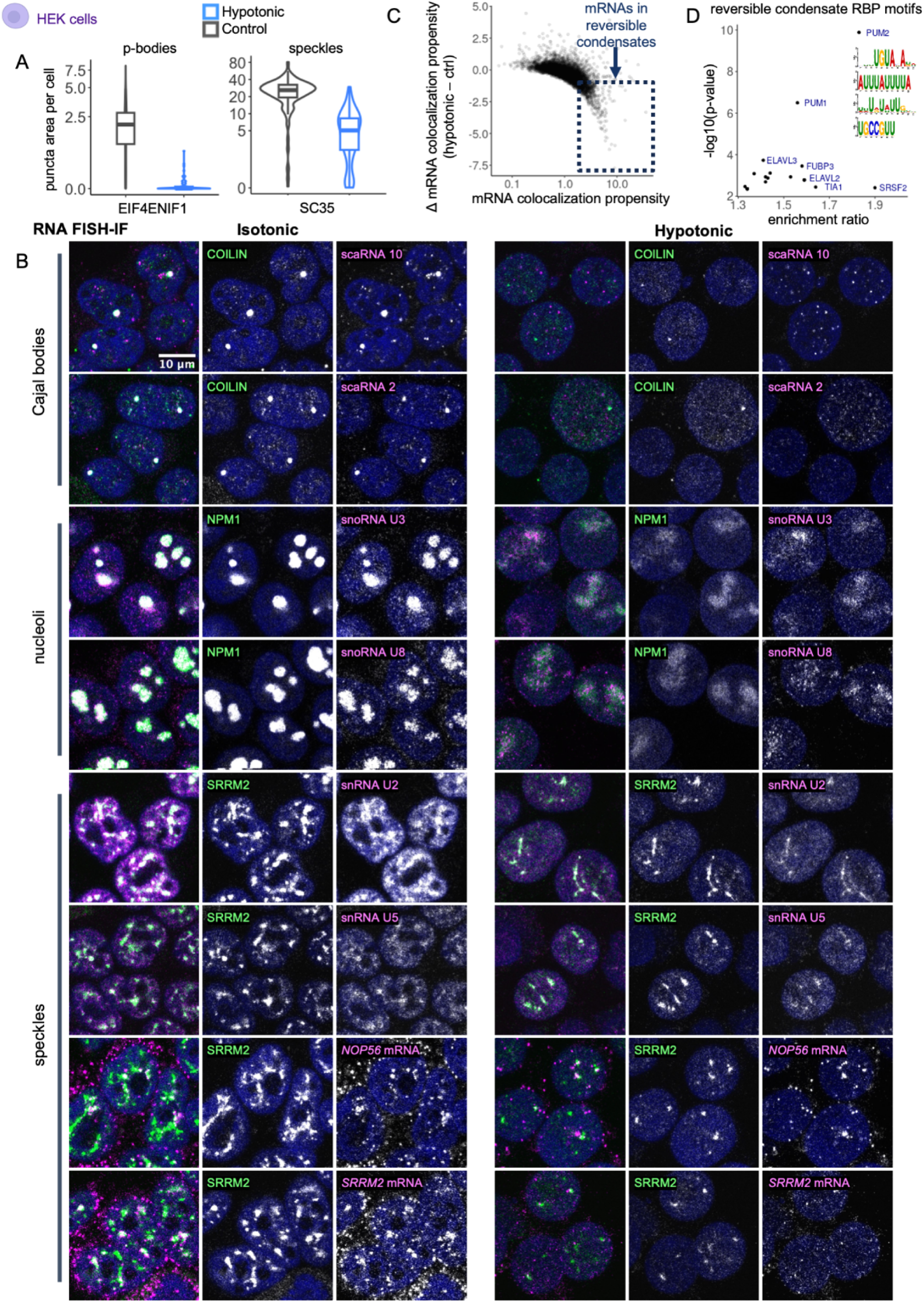
Hypotonic treatment uncovers reversible RNA colocalization, related to. Figure 2 (A) IF was used to measure total puncta area per HEK cell before and after hypotonic treatment (images in Figure 2B). Data presented as Tukey box plots with violin plots to show distribution and with arcsinh transform. (B) FISH-IF images of RNA enrichment in condensates before and after hypotonic treatment with nuclei stained with Hoechst, colored dark blue (quantification in Figure 2E and 2F). (C) mRNAs plotted, with those putatively enriched in reversible condensates identified by top quartile colocalization propensity in untreated conditions and top quartile sensitivity to hypotonic treatment (decrease in colocalization propensity upon hypotonic treatment). X-axis is log transformed. (D) RBP motifs enriched in the 3’ UTRs of putative mRNAs in reversible condensates identified in (C) vs length-matched control 3’ UTRs.

**Figure S4.**
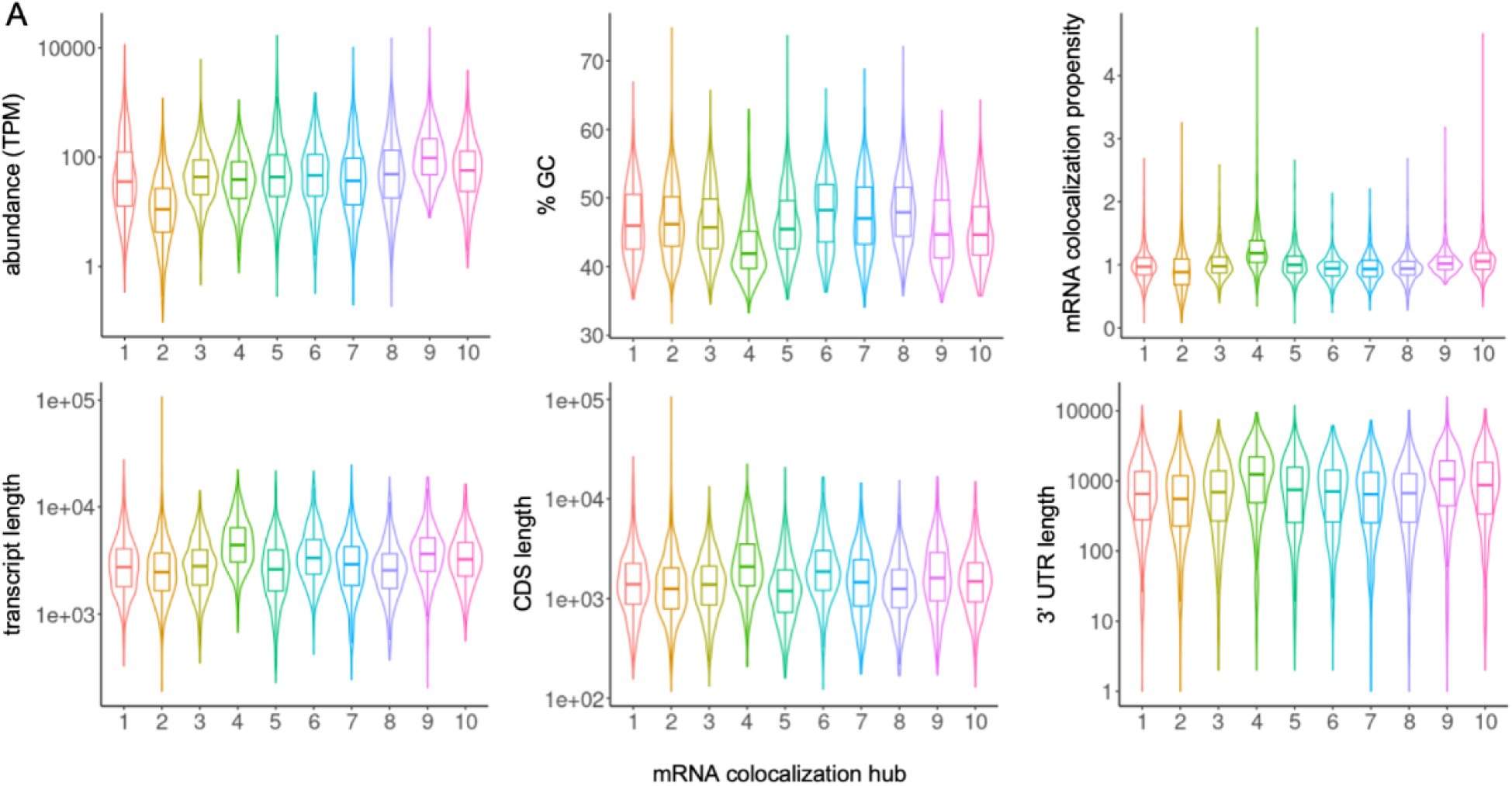
mRNA colocalization hub properties, related to. Figure 3 (A) Sequence features of mRNA colocalization hubs. Data presented as Tukey box plots with violin plots to show distribution. TPM and length axes are log transformed.

**Figure S5.**
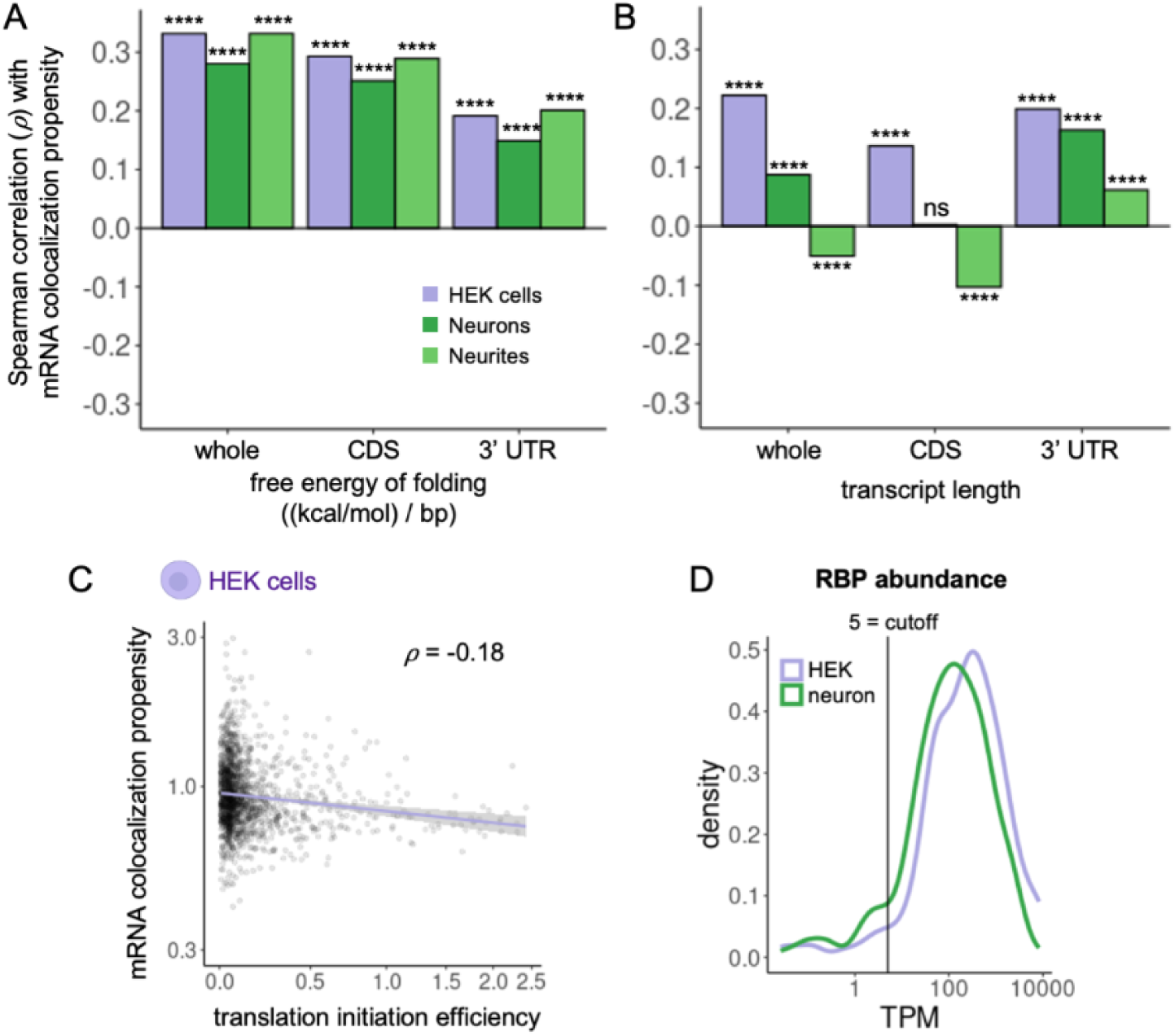
mRNA colocalization propensity correlations, related to. Figure 4 (A-B) The Spearman correlation of mRNA colocalization propensity vs. length normalized free energy of folding (A) or transcript length (B) for the whole transcript, the coding sequence (CDS), or 3’ UTR. (C) Individual mRNAs plotted to show correlation of mRNA colocalization propensity in HEK cells vs. translation initiation efficiency^104^ with Spearman correlation ⍴. X-axis is arcsinh transformed and y-axis is log transformed. (D) The distribution of TPM for the mRNAs encoding each RBP with an annotated motif^97,98^ is plotted for the HEK and whole neuron RNA-SPRITE datasets. A cutoff of 5 TPM was used for RBP motif analyses. \*\*\*\**p* < 0.0001

**Figure S6.**
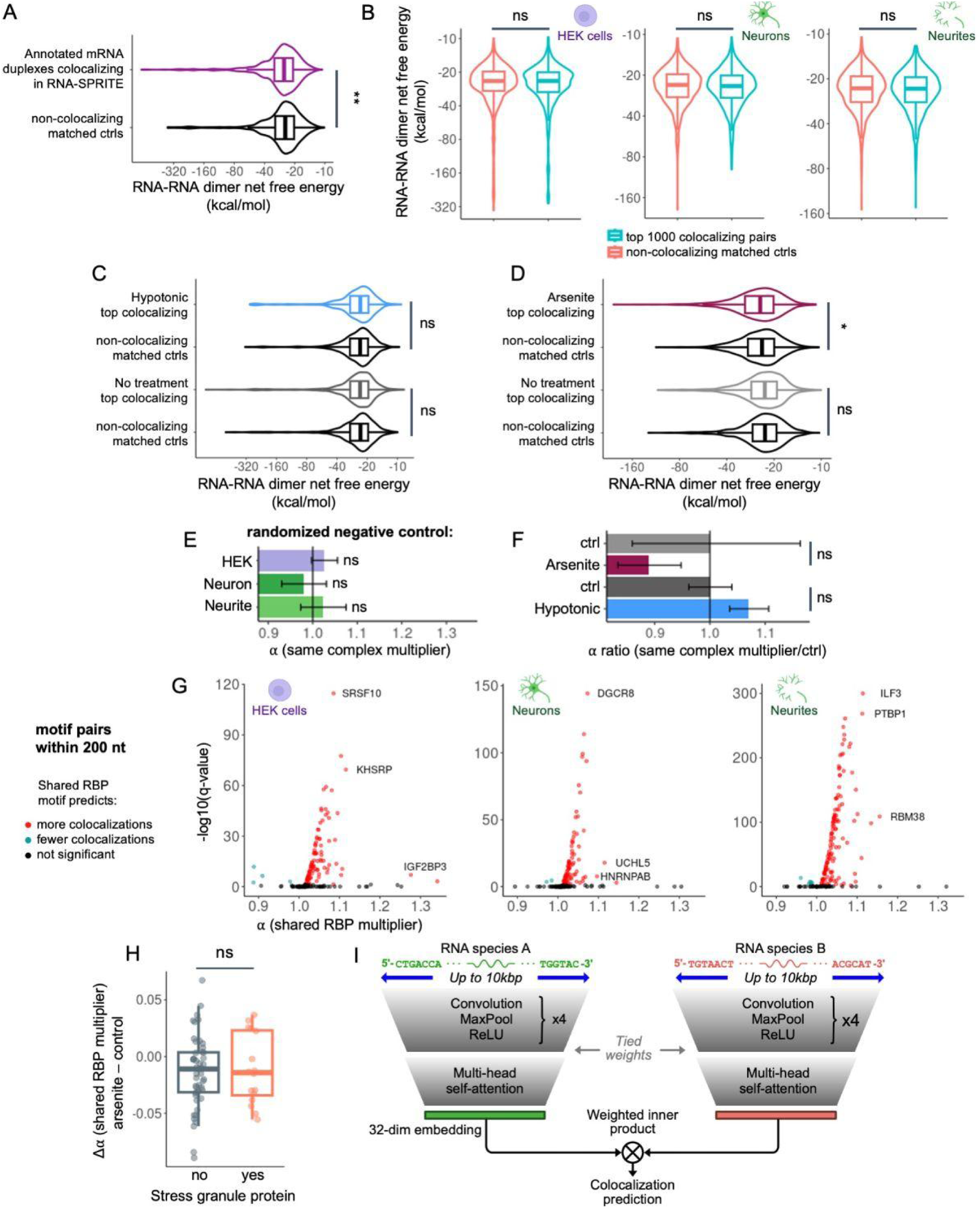
Drivers of mRNA colocalization pairwise specificity, related to. Figure 5 (A-D) Free energy of folding for intermolecular dimers minus values for separate monomers. Data presented as Tukey box plots with violin plots to show distribution and with arcsinh transform of y-axis. (A) mRNA pairs previously annotated to form a duplex^107^ that also colocalize in the HEK cell RNA-SPRITE data vs. non-colocalizing, non-duplex length-matched controls. (B-D) Highest confidence (by z-score) 1000 colocalizing mRNA pairs in each dataset vs. non-colocalizing length-matched controls. (E) As a negative control, the proteins annotated as interacting within a complex were randomly assorted to complexes, retaining the original number of constituents for each complex, number of complexes, and number of complexes each protein appeared in. The multiplicative effect on mRNA pairwise specificity (α) if two mRNAs encode proteins annotated to be in the same scrambled control complex. (F) The multiplicative effect on mRNA pairwise specificity (α) if two mRNAs encode proteins annotated to physically interact in the same complex, divided by the control condition. (G) For each RBP, the multiplicative effect on mRNA pairwise specificity (α) if two mRNAs are called as targets of the same RBP vs. Benjamini-Hochberg adjusted significance. An mRNA was considered a target of an RBP if it had two significant motifs for the RBP within 200 nucleotides anywhere in the transcript. (H) For stress granule enriched and non-enriched RBPs^117^, the change in the multiplicative effect on mRNA pairwise specificity (Δα) for arsenite treatment minus control. Data presented as Tukey box plot. (I) Neural network architecture for predicting colocalization pairwise specificity. (A-D, H) Wilcoxon rank-sum test (E-F) Error bars represent 95% confidence intervals. \**p* < 0.05, \*\**p* < 0.01, ns = not significant

## METHODS

### Cell culture and drug treatments

All cells were grown at 37°C with 5% CO2. HEK293T (HEK, ATCC) were grown with DMEM (GIBCO, 11995065), 10% FBS (Atlanta Biological, S11150H), and 1% penicillin-streptomycin (GIBCO, 15140122), according to standard protocols. For arsenite treatment conditions, neurons were treated with 0.5 mM sodium arsenite for 1 hour. For hypotonic treatments with HEK cells, 3 volumes of water were added to 1 volume of media for 10 minutes (min) before fixation. Fixation was performed as described below for each technique using diluted buffer (3 volumes of water to 1 volume of buffer). Conditions after fixation were the same as control.

### Primary cortical neuron culture

Pre-treated poly-D-lysine plates (Corning BioCoat) plates were used for whole neuron sequencing experiments. For neurite isolation experiments, PET millicell hanging cell culture insert, 1 µm pore size, in 6 well plates were treated with 0.01 mg/mL poly-D-lysine at 37°C overnight and were washed x4 in HBSS. Were used for neurite isolation. For imaging experiments, the inner 60 wells of 96 well glass bottom plates were treated with poly-D-lysine in the same way, with the outer 36 wells of the 96 well plate filled with ultrapure water. 6 mL (10 cm plate), 3.5 mL (hanging filters per well), 50 µL (96 well plate per well) of neuron media (Gibco Neurobasal Plus with 2% Gibco B27 Plus, 1% penicillin-streptomycin, and 250 ng/mL Amphotericin B) with 2% (1.14% for hanging filter wells) Gibco CultureOne supplement (antimitotic) was added to each well to keep the growing surface submerged, and the plates were stored at 37°C overnight, 5% CO2.

Embryos were collected from euthanized Sprague-Dawley rats (Charles River Laboratories) at embryonic day 17 via cesarean section. The embryos were transferred to HBSS in 10 cm glass plates, and were kept on ice to make the meninges easier to remove. The placenta was cut from each embryo, the heads were removed and transferred to a new glass plate with HBSS. Using a dissection microscope, the skull was removed and brains were transferred to a new glass plate with HBSS. Meninges were carefully and thoroughly removed, and the cortex was cut away from the striatum and other structures, and were transferred to a 10 mL conical with HBSS.

Worthington papain dissociation kit was used to dissociate cortices into individual cells in a biosafety cabinet using sterile technique. Reagents were prepared as described by the kit. HBSS was carefully removed from the cortices and 5 mL papain solution was added (100 units papain, 1000 units DNase I, 1mM L-cysteine, 0.5mM EDTA in HBSS). The conical was inverted x3 then incubated at 37°C for 20 minutes, with no agitation or inversion after the incubation. The papain solution was removed, and 3 mL of inhibitor solution (3 mg ovomucoid inhibitor, 3 mg albumin, and 500 units DNase I in HBSS) was added to the cortices, inverted x3, and sat upright for 5 min. Supernatant was removed and replaced with 3 mL additional inhibitor solution, inverted x3, and sat upright for 5 min. Supernatant was removed and 1.5 mL neuron media was added. A flame-treated Pasteur pipette was used to slowly triturate up and down 10x, avoiding bubbles. Cells were allowed to settle in the upright tube for 2 min. The top 750 µL of dissociated cells were removed and added to a new 10 mL conical. 750 µL neuron media was added to the original tube, triturated 10x, settled in the upright tube for 2 min, and the top 750 µL of dissociated cells were transferred to the new 10 mL conical. This process was repeated 1 more time, for a total of 3 trituration steps, adding all of the media with cells to the new tube after the final trituration. Cells were centrifuged for 5 min at 300 g, supernatant was removed, resuspended in 1 mL neuron media, and counted using a hemocytometer.

Cells were diluted in additional neuron media to achieve 6 million cells in 6 mL for 10 cm plates, 0.5 million cells per hanging filter in 0.5 mL, 25,600 cells in 50 µL per well 96 well plates. The evenly-suspended, diluted cells in the indicated media amounts were added to each well of the previously prepared plates to bring the volume to 12 mL for 10 cm plates, 4 mL for hanging filters, and 100 µL for 96 well plates each with 1% CultureOne supplement final. CultureOne supplement was not used again after this treatment on day in vitro (DIV) 0. On DIV3 100 µL more neuron media was added to the 96 well plate to bring the final volume to 200 µL. Twice per week (96 wells or hanging filters) or three times per week (10 cm plates), half of the media was removed from each well and replaced with fresh media and plus ultra-pure water to counter evaporation. Experiments were performed on DIV17.

### Immunofluorescence (IF)

Cells were fixed with freshly mixed 4% paraformaldehyde (PFA) in phosphate buffered saline (PBS) for 10 min to fix. Washed x2 in Tris-buffered saline with 0.1% Tween-20 (TBST). Cells were permeabilized with TBST with 0.1% Triton-X for 15 min at room temperature (RT), washed 2x TBST, and blocked with TBST with 10% normal goat serum for 1 hr RT. Cells were treated with primary antibodies and 10% normal goat serum overnight, washed 3 times for 5 min each, and then treated with secondary antibodies diluted in TBST with 10% normal goat serum for 2-3 hrs RT. Cells were then washed 3 times for 5 min each. If needed, Hoechst was added to the second to last wash (1:5000) or DRAQ5 (1:5000) with no wash out.

### Fluorescence in situ hybridization (FISH)-IF

SABER FISH was performed according to prior protocols^139^ in 96 well plates using PaintSHOP^140^ for probe design, with 21-30 oligos per mRNA. For target probe incubation, 0.5 ug each of two concatemerized target probes in 50 uL total solution volume overnight at 43°C was used. Fluorescent imaging probes were used at 0.2 uM in 50 uL total solution volume, incubated at 37°C for 15 min. IF was performed after FISH steps. Cells were blocked for 1 hr at RT in blocking buffer: PBST (1x PBS with 0.1% Tween-20) with 10% molecular grade RNase free Bovine Serum Albumin and 1:100 murine RNase inhibitor (NEB). Cells were incubated overnight at 4°C with primary antibodies in blocking buffer, washed with PBST twice for 10 min each, and then treated with secondary antibodies for 1-2 hrs at RT in blocking buffer. Cells were washed with PBST twice for 10 min each then stored in PBST with 1:100 murine RNase inhibitor at 4°C or immediately imaged.

### Plasmids, lentiviral packaging, and transduction

FM5 mGFP-DDX6, FM5 mGFP-Dcp1a, and FM5 mGFP-G3BP were a kind gift of David W. Sanders. FM5 NPM1-mCherry was a kind gift of Lian Zhu. FM5 SFFV Halo-FMR1 was cloned using NEBuilder HiFi DNA Assembly Cloning Kit (NEB E5520) and the FM5 lentiviral backbone vector (a gift from David W. Sanders)^91^. Lentivirus was made using typical protocols for third generation packaging vectors and concentrated with Lenti-X concentrator (Takara Bio 631231). pHAGE RSV GCaMP6s (addgene #80146) was used for imaging activity of cultured neurons.

### Imaging and analysis

A Zeiss 980 laser scanning confocal microscope was used for imaging with a 60x oil objective unless otherwise noted. GCaMP6 activity videos were taken with a 10x air objective. Image analysis was done using custom Fiji macros and R scripts. For puncta area per cell measurements, whole-cell z-stacks were taken, and maximal intensity projections were used for analysis after a one pixel Gaussian blur to de-noise. A single threshold was chosen for each condensate marker protein based on a range of images, and condensate puncta with area 0.01 to 2 um^2^ were quantified. A custom Fiji macro was used to measure the total area of puncta for each image and then divided by the number of cells per image (quantified by counting nuclei with a nuclear marker Hoechst or DRAQ5).

For partition coefficient analysis, nuclei were segmented using Hoechst signal after a two pixel Gaussian blur, and nuclear bodies were segmented using the indicated condensate markers after a one pixel Guassian blur. A single segmentation threshold was used for each condensate marker based on a range of images, and condensate puncta with area 0.01 to 2 um^2^ were quantified. For each nuclei, Fiji was used to measure the fluorescence intensity inside the segmented nuclear condensates and outside the condensates in the nucleoplasm was measured. The partition coefficient is the ratio of the mean fluorescence intensity inside the segmented condensate divided by the mean fluorescence intensity outside of the condensates in the nucleoplasm, measured per cell.

For FISH validation of RNA-SPRITE Louvain colocalization hubs, 6-7 mRNAs per hub were chosen and each given 1 of 3 unique hairpin sequences. Colocalization was quantified for every possible mRNA pair, as long as they did not share the same hairpin sequence and were therefore compatible for multiplexing. The rodent primary neurons were imaged at a plane just under the nucleus. UPF1 IF was used to mark the cell body cytoplasm and segmented using 10 pixel Gaussian blur and a consistent mean fluorescence threshold for segmenting. MAP2 IF was used to mark dendrites and segmented using 2 pixel Gaussian blur and a consistent mean fluorescence threshold for segmenting. Two mRNA targets (visualized using FISH as above) were imaged per well in separate fluorescence channels. For both FISH channels, a one pixel Guassian blur was done to de-noise. The first FISH channel was segmented using a consistent mean fluorescence threshold and filtering for 0.1 um^2^ minimum FISH puncta area. The fluorescence intensity of the second FISH channel and the UPF1 and MAP2 channels was measured. Dendrite vs. cell body puncta were identified by their relative fluorescence intensity in the UPF1 and MAP2 channels.

A single FISH puncta in the first channel was considered positive for colocalization of the two mRNAs if the measured FISH channel maximum fluorescence value was above a consistent threshold. The same process was repeated for the second FISH channel. The percent positive puncta for each thresholded FISH target and measured FISH target pair was calculated (each mRNA pair would have two values, one based on thresholding on in the first FISH channel and one based on thresholding in the second FISH channel). For each measured mRNA target, the median percent positive puncta for mRNA pairs not in the same RNA-SPRITE Louvain colocalization hub was calculated. The percent positive puncta for all mRNA pairs was divided by this number to give “single normalized colocalization,” which is normalized for the abundance of measured mRNA. For each thresholded mRNA target, the median “single normalized colocalization” for mRNA pairs not in the same RNA-SPRITE Louvain colocalization hub was calculated. All “single normalized colocalization” values were divided by this number to give “double normalized colocalization” in order to normalize for the abundance of the thresholded mRNA target in addition to normalizing for the abundance of measured mRNA target. “Double normalized colocalization” is plotted in Figure 3D and 3E for mRNAs in the same hub. Values over 1 indicate within hub specificity.

Normalized colocalization for mRNAs encoding proteins that physically interact was performed in the same way, but using abundance matched control mRNAs instead of mRNAs in a different colocalization hub. The proteins encoded by *Apc* and *Ctnnb1* physically interact. *Picalm* was the matched control for *Apc* and *Tpm1* was the matched control for *Ctnnb1*.

### RNA-SPRITE crosslinking

Low passage HEK cells were plated on pre-treated poly-D-lysine plates. Neurons were prepared as above. HEK and whole neurons were crosslinked in 10cm plates. Neurites were crosslinked in 6 well hanging filters. Crosslinked was performed according to the following protocol with RNase-free reagents.

One bottle of phosphate-buffered saline (PBS) was chilled at 4°C. One was kept at room temperature (RT). A 2mM Disuccinimidyl glutarate (DSG) stock solution was prepared in room-temperature PBS: DSG was allowed to sit for 20 min at RT to avoid moisture, then 306 µL DMSO was added to a 50 mg bottle of DSG (0.5M final concentration). The solution was vortexed to mix. It was used fresh or within 1 month.

Media was removed from plates. From this point, all RNA-SPRITE steps were performed using certified RNase-free reagents and consumables wherever possible, and RNaseZap was used on gloves, equipment, and all surfaces. Cells were washed with room-temperature 1 volume (5 mL for a 10 cm plate, 2 mL for hanging filters in a 6-well plate) 1x PBS. PBS was gently removed. One volume of 2mM DSG solution was added. The plates were rocked gently at room temperature for 45 minutes. During this time, RT and chilled 1x PBS were prepared in a 50 mL conical tube, 1 falcon tube, and 1 microcentrifuge tube per sample. Right before use, in a fume hood, a 1% formaldehyde solution was premixed in room-temperature PBS using a fresh ampule. The DSG solution was removed from cells, and the cells were washed once with room-temperature PBS. One volume of formaldehyde solution was added to the cells. The cells were incubated at room temperature for exactly 10 minutes, gently rocked or swirled by hand occasionally. Immediately, 200 µL of 2.5M Glycine stop solution per 1 mL of formaldehyde solution in the dish was added. The mixture was incubated at room temperature for 5 minutes, shaken or gently swirled by hand occasionally. Crosslinking reagents were poured off into formaldehyde waste. The cells were carefully washed with cold 1x PBS, gently rocked for 1-2 minutes. The wash step was repeated 1-2 more times, discarding into formaldehyde liquid waste.

After the last wash, one volume of scraping buffer (ice-cold PBS + 0.5% bovine serum albumin) was added to each well/plate. From this point onward, the cells were kept at 4°C. The cells were scraped from the plate (underside of filters for neurite collection) and transferred to a 15 mL falcon tube. Each well was rinsed with another volume of scraping buffer, which was added to the falcon tube. The cells were centrifuged at 1000 g at 4°C for 5 min to pellet them, and the supernatant was discarded. The pellet was resuspended in 1 mL cold scraping buffer to break it up. The cells were aliquoted into microcentrifuge tubes and spun at 2000g for 5 min at 4°C. The supernatant was removed, and the cells were flash frozen in liquid nitrogen.

### RNA-SPRITE Lysis

A Roche complete EDTA-free protease inhibitor tablet was dissolved in 500 µL (1 tablet per 10 mL). Buffers were prepared and used within 1 month: lysis buffer A (50 mM Hepes pH 7.4, 1 mM EDTA, 1 mM EGTA, 140 mM NaCl, 0.25% Triton-X, 0.5% NP-40, 10% Glycerol, Ultra Pure H2O), lysis buffer B (50 mM Hepes pH 7.4, 1.5 mM EDTA, 1.5 mM EGTA, 200 mM NaCl, Ultra Pure H2O), lysis buffer C (50 mM Hepes pH 7.4, 1.5 mM EDTA, 1.5 mM EGTA, 100 mM NaCl, 0.1% Na-DOC, 0.5% NLS, Ultra Pure H2O). Lysis buffers A, B, and C were chilled on ice. The centrifuge was chilled, and all steps were performed on ice.

A cell pellet was thawed on ice for two minutes or until thawed. To each pellet, 700 µL of Lysis Buffer A supplemented with 1x protease inhibitor + 1:40 murine RNase inhibitor (NEB) was added, and the pellet was fully resuspended by pipetting. It was ensured that the pellet was fully resuspended in the lysis buffer. The mixtures were incubated on ice for 10 minutes. They were not inverted or mixed after incubation. The cells were pelleted at 4°C for 8 minutes at 850 × g. The supernatant was discarded, taking care not to disturb the pellet (some buffer was left rather than disturbing the pellet). To each cell pellet, 700 µL of Lysis Buffer B supplemented with 1x protease inhibitor + 1:40 RNase inhibitor was added, and the pellet was fully resuspended. The mixtures were incubated on ice for 10 minutes without inversion or mixing. The cells were pelleted at 4°C for 8 minutes at 850 × g. The supernatant was discarded, taking care not to disturb the pellet. To each 10 million nuclei pellet, 1.1 mL of Lysis Buffer C supplemented with 1x protease inhibitor + 1:40 RNase inhibitor was added, and the pellet was resuspended. The mixture was incubated on ice for 8 minutes. Some lysate was taken to check RNA sizes pre-sonication. The lysate was flash frozen and stored at −80°C or used for sonication.

### RNA-SPRITE sonication and checking RNA size

1 mL of sample was transferred to a 1 mL AFA tube. Covaris ME220 sonicator instructions were followed with the following parameters: 9°C, 6-12°C range, 75 PIP, 1000 CPB, 15 duty factor, 2 min. Single-use aliquots were made and flash frozen, then stored at −80°C.

The crosslinks were reversed on a 20 µL aliquot of each sample before and after sonication by adding 25 µL NLS elution buffer (20 mM Tris-HCl pH 7.5, 0.56 M NaCl, 10 mM EDTA pH 8, 2% N-lauroylsarcosine, Ultra Pure H2O) and 4 units of Proteinase K (NEB), then the mixture was shaken at 1200 rpm on a thermomixer for 1 hr at 65°C. The samples were cleaned up by following the protocol provided in the Zymo RNA Clean and Concentrator-5 Kit (>17 nt kit). DNase digestion was done with 41.75 µL sample, 1x Turbo DNase buffer, 4 units Turbo DNase, and 1:40 murine RNase inhibitor. The mixture was incubated for 30 min at 37°C. Samples were cleaned up with the Zymo RNA Clean and Concentrator-5 Kit and eluted in 14 µL.

A Qubit (HS RNA) and Tapestation (HS RNA) were used to measure RNA concentration and size according to the manufacturer’s instructions. The RNA size range should be similar before and after sonication, and the RNA integrity number should be over 5 for both. Sonication time and/or duty factor might need to be decreased if RNA is more degraded.

### RNA-SPRITE NHS bead coupling

We calculated the volume of crosslinked sample to add per volume of beads to achieve 5 molecules of RNA per bead using estimates from Qubit and Tapestation data on reverse crosslinked samples above. We used 4 mL total of NHS beads per experiment, dividing the beads equally between each sample in 1.5 mL DNA lo-bind tubes. The volumes below are for an experiment with 10 samples (including replicates) that were kept separate until split pool barcoding. If the amount of RNA per tube was changed (e.g. by using a different number of samples), the volumes per tube would need to be changed proportionally for all reactions before the split pool barcoding. The samples could not be frozen again until the step where they were eluted from the beads by Proteinase K and reverse crosslinking treatment. Therefore, we performed all steps from bead coupling to bead elution on 4 consecutive days for each experiment. The mixing steps were done with a thermomixer.

The bottle containing the Pierce NHS-activated beads in N,N-dimethylacetamide (DMAC) was vortexed until a uniform suspension was obtained. The time the bottle was open was minimized, the lid was sealed with parafilm, and the DMAC bead tube was stored in a 50 mL conical with drierite at 4°C. A total of 4 mL was transferred into clean 1.7 mL protein lo-bind tubes, with a maximum of 1.2 mL per tube. The tube was placed on a magnetic rack to capture the beads (the magnetic rack was used for all subsequent washes in the RNA-SPRITE protocol as well). The DMAC was removed, and the beads were washed with 1 mL ice-cold 1 mM HCl (from a 1 M HCl stock). The beads were washed with 1 mL ice-cold 1x PBS. 500 µL of RT Coupling Buffer (1x PBS, 0.1% SDS, 5 mM EDTA) with 1:100 NEB murine RNase inhibitor was added to the beads, vortexed, and was not put on ice. 1:10 dilutions of each lysate were made for NHS coupling if necessary. It was ensured that samples were never spun down prior to bead coupling. The mixture was vortexed heavily to ensure all material was in solution prior to coupling. Lysate was added to RT Coupling Buffer with 1:100 RNase inhibitor to a final volume of 500 µL, and the mixture was vortexed well to ensure it was evenly mixed. Then, 500 µL of beads was added to each sample and vortexed well with beads. The lysate and beads were incubated at room temperature for 15 min, then overnight at 4°C on a rotator.

### RNA-SPRITE RPM ligation

The NHS beads were quenched and washed by placing tubes on a magnet and removing half of the total volume of flowthrough (500 µL). 500 µL of 1M Tris-HCl pH 7.5 was added to the beads, vortexed, and incubated at RT for at least 45 minutes, 1000 rpm on the rotator. This ensured that all NHS beads were quenched with protein from bound lysate or Tris and would not bind enzymes in the following steps. Beads were washed three times in RT RLT++ Buffer, and the lid was rinsed. Beads were washed 4x times with RT M2 Buffer (20mM Tris-HCl pH 7.5, 50mM NaCl, 0.2% Triton-X, 0.2% NP-40, 0.2% Na-DOC, Ultra Pure H2O), and the lid was rinsed (0.5 to 1 mL washes were used from here on unless otherwise stated). On ice, 100 µL of PNK master mix (1x PNK buffer, 80 units T4 PNK Enzyme, 1:40 murine RNase inhibitor, Ultrapure water) was added to each sample to convert to 3’OH RNA ends. The mixture was incubated at 37°C for 45 minutes, shaking at 1200 rpm on the thermomixer. 1x wash in RLT++ buffer (1 x Buffer RLT, 10mM Tris-HCl pH 7.5, 1mM EDTA, 0.2% NLS, 0.1% Triton-X, 0.1% NP-40) was performed. 4x washes were performed in M2 Buffer.

The RPM ligation reaction (7.5% DMSO, 5 µM RPM adapter, 1x NEB T4 RNA Ligase Buffer, 1 mM ATP, 15% PEG 800, 1:200 murine RNase inhibitor, and 120 units NEB T4 RNA Ligase High Concentration) was prepared to a total volume of 100 µL per sample. The water, DMSO, and RPM adapter were added to the beads first, heated at 65°C for 2 minutes at 1000 rpm to denature the secondary structure of RNA, and then immediately placed on ice for at least 3 minutes. The tubes were vortexed and touch spun 2x. The rest of the RPM ligation reaction components were added, and the tubes were flicked to mix. The mixture was mixed well by vortexing and very brief pop-spins repeatedly after adding the ligase. Care was taken to ensure there wasn’t a bead clump on the bottom. The mixture was incubated for 1 hour at 24°C at 1200 rpm with mixing. Beads were washed 4x in No Salt Tween/NP40 Buffer (50mM HEPES pH 7.4, 0.1% Tween, 0.1% NP-40, Ultrapure H2O) to remove excess RPM adapter. 1x wash with RLT++ buffer and 6x washes with M2 buffer were performed.

Residual solution was completely removed from NHS beads. The reverse transcription mastermix (0.25 µM RPM bottom adapter, 1x First Strand buffer, 5 mM DTT, 0.5 mM dNTPs, 500 units Superscript III enzyme, 1:40 murine RNase inhibitor) was prepared to a total volume of 100 µL per sample. The RPM bottom oligo and water were added first. The mixture was heated at 65°C for 3 minutes at 1000 rpm to denature the secondary structure of RNA, and then immediately placed on ice for 3 minutes. The tubes were vortexed and touch spun 2x. Vortexing was performed to make sure beads were fully resuspended prior to adding the RT mastermix with enzyme, or mixing was done by pipetting after the full mastermix was added. The rest of the reverse transcription master mix components were added on ice to prevent mispriming. The mixture was incubated at 42°C for 30 minutes, 1200 rpm. Beads were washed 4x with M2 Buffer. Residual solution was completely removed from NHS beads. To digest excess RPM bottom adapter, beads were resuspended in 100 µL Exo1 master mix (1x first strand buffer, 1:40 murine RNase inhibitor, 200 units NEB Exonuclease I, and ultrapure water) and incubated at 37°C for 20 minutes, 1200 rpm. Excess RPM was washed 6x with M2 Buffer, ensuring that tube lids were washed out. Beads could be stored at 4°C overnight in 0.25-1 mL M2 Buffer with 1:40 dilution of Murine RNase Inhibitor and 5 mM EDTA to prevent RNA degradation overnight.

### RNA-SPRITE Split-pool barcoding

The process for split-pool barcoding was prepared by washing 3x with M2 with rinsing out the lid. 4 total rounds of adapter ligations were performed: Odd, Even, Odd, and Terminal Y. 96 adaptors were designed to ligate onto the RNA molecules. The ligation reaction between the adaptors and the RNA occurred in a 96-well plate. Efforts were made to lose as few beads as possible with each step to maximize yield. A plate holder was always used for 96-well plates.

SPRITE barcode plates were prepared by diluting 1:10 to 0.45 μM final concentration from a 4.5 μM stock in 1x Annealing Buffer. The SPRITE adaptor stock plate was centrifuged at 1000 x g for 1 minute before the foil seal was removed. The dilution was performed by adding 21.6 µL of 1x annealing buffer to the existing 2.4 µL of 4.5 μM plate, yielding a 24.0 µL final volume for a 1:10 dilution. The mixture was mixed well by pipetting up and down, and a foil seal was added. The plate was spun down at 1000 x g for 1 minute before the foil seal was removed. 2.4 µL was aliquoted from the new 0.45 μM plate of SPRITE adaptors to a new low-bind 96-well plate. Care was taken to ensure that there was no mixing between wells at any point of the process to avoid cross-contamination of barcodes. A new pipette tip was used for each well. After the transfer was completed, both plates were sealed with a new foil seal. 3.5 mL of Ligation Master Mix (2.5x NEBNext Quick Ligation Reaction Buffer, 0.63x Instant Sticky-end Ligation Master Mix, 19% 1,2-Propanediol) was made for four rounds of SPRITE, and the remainder was stored at −20°C. The master mix was split evenly into each well of a 12-well strip tube by pipetting ∼240 µL into each tube (6.4 MM per well x 8 wells x 4 plates). The master mix was kept on ice. A dilute M2 wash buffer was created by mixing M2 wash buffer with an equal volume of H2O and 1:200 murine RNase Inhibitor to resuspend beads for each round. The samples were kept separate for the first round of barcoding. Wells for each sample were recorded in round one for later identification.

To begin each round, ∼955 µL of dilute M2 wash buffer was added to the beads to achieve a final volume of 1.075 mL total for all samples. Care was taken to ensure that the beads were equally resuspended in the buffer. The samples were transferred into a strip tube for easy pipetting into the plate (1 strip tube for a plate). For subsequent rounds when all the samples were mixed, the beads were distributed equally into a 12-well strip tube by aliquoting ∼92 µL of beads into each well.

The 96-well plate containing the aliquoted adaptors was centrifuged at 1000 x g for 1 minute, and then the foil seal was removed. 11.2 µL of beads was aliquoted into each well of the 96-well plate that contained 2.4 µL of the 0.45 μM SPRITE adaptors. Care was taken to ensure that there was no mixing between wells at any point of the process. A new pipette tip was used for each well. Care was also taken to ensure that there were no beads remaining in the pipette tip or on the walls of the well. Any remaining beads were carefully added to individual wells on the plate in 1 µL aliquots. 6.4 µL of Ligation Master Mix was aliquoted into each well, and the mixture was allowed to mix in solution by placing the plate on a mixer at 1600 rpm to avoid loss on tips. The plate was sealed very well with a foil seal, rolling in a grid pattern, and incubated on a thermomixer for 30-60 minutes at 20°C, shaking for 2 minutes at 1600 rpm. Wells were checked to ensure they were well mixed, then the plate was shaken at 1600 rpm for 5 minutes, checked, and then at 1200 rpm continuously to prevent beads from settling to the bottom of the plate.

Each 20 µL ligation reaction was quenched by transferring 60 µL of M2 SPRITE Wash buffer + 50 mM EDTA from a sterile plastic reservoir into each well of the plate. The mixture was incubated for 1-2 minutes before pooling. Tips were not touched to the inside of wells if reused. All 96 stopped ligation reactions were pooled into a new sterile plastic reservoir. From this point in the first round of barcoding, all samples were mixed. Each well was rinsed with additional M2 + EDTA to collect all beads. A 15 mL conical tube was placed on an appropriately sized magnetic rack, and the ligation pool was transferred into the conical. The beads were captured on the magnet. The reservoir and serological pipette were rinsed with cleared supernatant repeatedly until fully clean to collect all beads. The cleared supernatant was discarded. The 15 mL conical containing the beads was removed from the magnet, and the beads were resuspended in 1 mL SPRITE Wash Buffer. The bead solution was transferred into 2 microcentrifuge tubes. Another 1 mL of wash buffer was used to rinse the conical, and this solution was transferred into the microcentrifuge tubes. The cleared supernatant was used to clean the conical until fully clean. The beads were washed 4x with 1 mL of SPRITE Wash Buffer. Care was taken to wash out the lids, and the tubes were spun after removing the washes to remove all liquid from each wash. The tubes were combined in the last wash, and an extra tube was rinsed with cleared supernatant. The beads were spun down briefly in a microcentrifuge and placed back on the magnet to remove any remaining liquid. The steps starting from “beginning of each round” were repeated for a total of four SPRITE rounds (Odd, Even, Odd, Terminal Y).

### RNA-SPRITE reverse crosslinking

The beads were resuspended in 1.275 mL NLS Elution Buffer. The sample was vortexed well and split evenly into aliquots in PCR strip tubes (48 wells total). Care was taken not to transfer any material between aliquots between this point and PCR. 2 units of proteinase K (NEB P8107S) in 10 µL NLS elution buffer were added to each reaction. The reaction was incubated at 50°C overnight (minimum 12 hrs) to reverse crosslink. The incubation was performed with continuous shaking at 1200 rpm to prevent beads from settling to the bottom. The tubes were taped down really well in two directions to make sure the lids did not come open and the tube strips stayed in the mixer. A thermolid was used to help avoid condensation on the lids. Care was taken to ensure the lids were sealed. The beads were vortexed, touch spun, placed on a magnet, and 30 µL was transferred to a PCR strip tube. The beads were rinsed in an additional 10 µL 1x PBS, vortexed, and spun down. The two eluates were combined to 40 µL total. The eluates could be frozen for later processing at this point or moved to the next step.

### RNA-SPRITE second reverse transcription and splint ligation

The silane beads were taken out (576 µL total) and placed in a 1.7 mL tube. The beads were separated using a magnet, and the buffer was discarded. The beads were washed 1x with Qiagen RLT buffer. The beads were resuspended in 1.2 mL RLT + 0.1% Triton-X (3x volume). 95 µL RLT + 0.1% Triton-X was added to each microcentrifuge tube, and then 25 µL beads in RLT + 0.1% Triton-X were added to each tube. The tubes were briefly vortexed (ensuring all lids were held closed) or mixed well by pipetting, and the liquid was brought to the bottom of the tube and touched spun. The samples were incubated for 1 minute at RT. 100 µL 100% EtOH (0.625x combined volume) was added to each tube and pipetted to mix. The samples were incubated for 5 minutes at RT to bind. The tubes were placed on a magnet, and the beads were moved back and forth to form a single clump before the supernatant was discarded. The beads were washed 3x with 150 µL 80% EtOH, keeping the beads on the magnet to keep them clumped, and rotated 6x. For one strip of tubes at a time, immediately after removing the final ethanol wash, the tubes were touch spun, ensuring a nice bead clump, and the remainder was removed with a P20. The beads were dried at RT until no longer shiny and eluted in 12.5 µL ddH2O, collecting 11.5 µL total in two rounds of elution. The samples were pipetted up and down and let sit for 2-5 minutes to elute.

A second round of reverse transcription was performed to complete the process after reverse crosslinking. To prevent promiscuous random priming, 4 µL of 5x First Strand Buffer was added ON TOP of the 11 µL sample, and then the sample was preheated at 40°C for 3 minutes (prior to adding Superscript III enzyme). While the sample was still at 40°C, 5 µL of RT mastermix (5 mM DTT, 1 mM dNTPs, 200 units Superscript III enzyme, 1:40 murine RNase inhibitor) was added directly on top of the 15 µL sample + First Strand buffer. The tubes were pipetted or flicked to mix, and the liquid was brought to the bottom of the tube with a touch spin. The samples were incubated at 40°C for 5 minutes, then 50°C for 30 minutes, followed by heat inactivation at 70°C for 15 minutes, and finally held at 4°C.

Separately, the splint was annealed together with 180 µM Splint Top-2puni-spcr, 180 µM Splint Bot-2puni-5phos-3spcr, and annealing buffer (10 mM Tris-HCl pH 7.5, 200 mM LiCl, Ultra Pure H2O). The annealing process was incubated at 95°C for 2 minutes, followed by 70 cycles of −1°C/20s (slow cool), and held at 10°C. The annealed splint was checked for dimers using a gel. 45 µM aliquots were made and diluted in annealing buffer, and then stored at −20°C. 2 µL of RNAseH + 1 µL RNase Cocktail was added directly on top of each sample to degrade RNA prior to cDNA ligation. The samples were incubated at 37°C for 30 minutes. 1 µL 45 µM splint and 3.2 µL water were added to 23 µL of the sample and mixed well. 12.8 µL Ligation Master Mix was added, and the tubes were mixed up and down 10 times. The samples were incubated at 20°C for 2 hours, shaking at 1200 rpm continuously.

The samples were cleaned with SPRI beads. The SPRI beads were brought to RT and vortexed. 32.5 µL of SPRI beads (0.8125x) was added to each sample. The samples were mixed well by pipetting and incubated at room temperature for 10 minutes (mixed by pipetting every 3 minutes). The samples were placed on a magnet for 3 minutes and the supernatant was discarded. The beads, still on the magnet, were washed 3x with 180 µL of 80% EtOH, 30 seconds each time. The tubes were lifted up halfway and tilted forward during liquid removal and addition to keep a nice single clump (rotation of tubes was not necessary). After removing the last wash, the remainder was removed with a P20. The beads were dried for 4-5 minutes at room temperature (until no longer shiny; the beads were observed to start cracking, but care was taken not to over-dry). The beads were eluted in 21 µL ddH2O total in two elutions. The samples were left to sit for 2-5 minutes both times to elute, recovering 20 µL total.

### RNA-SPRITE library prep and sequencing

PCR was performed with one strip tube of aliquots first to check if the number of cycles gave appropriate amplification (see below) before proceeding with the rest of the tubes. Each tube was assigned a unique i5/i7 index primer pair. PCR was performed with a total reaction volume of 50 µL per tube using 0.5 µM i5 index primer, 0.5 µM i7 index primer, and 1x Q5 Hotstart mastermix. The tubes were flicked to mix, and touched spun. PCR conditions were set as follows: (98°C 40s) 1 cycle, (98°C 15s; 69°C 15s; 72°C 90s) 4 cycles, (98°C 15s; 72°C 90s) 10 cycles, (72°C 2min) 1 cycle. The samples were cleaned with SPRI beads as described above, but with 37 µL SPRI beads (0.74x) per tube. The samples were eluted with 12 µL water total in two rounds of elution. The samples were run on a DNA HS D1000 TapeStation per the manufacturer’s instructions. We aimed for the 300-1000 bp region to be around 10 nM per tube, with 2 to 20 nM considered acceptable. The number of PCR cycles was adjusted for the second PCR set (up or down from 10 cycles) for the remaining tubes, if warranted.

To remove all primer dimer, the libraries were pooled and gel cutting and clean up were performed 1 to 2 times to extract the appropriate library sizes. Equal moles of libraries were pooled. No more than 100 ng, 20 µL per lane were loaded, using a 2% E Gel, with 20 µL water loaded into empty lanes and the ladder separated from the sample by 2 lanes. The gel was run for 8-13 minutes. The gel was gently cracked open, and a clean razor was used to cut. The gel was cut at ∼300-1000 bp. The primer dimer, at ∼250 bp, was observed just above the cut site. The Zymo gel cleanup kit was used according to the manufacturer’s instructions: the gel was melted thoroughly at 55°C for 5-10 minutes, a 1C column was used, two washes were performed, a 2-minute dry spin with a new tube was done, and the sample was eluted in 12 µL. The sample was diluted 1:10 for DS D1000 Tapestation to check for dimer. If any dimer was left, a second gel cut was performed.

For RNA-SPRITE Sequencing, Read1 had the RPM adapter and the RNA sequences to be aligned. Read2 had the barcode and required sequencing of at least 87 bp: (24 bp x 3 barcodes) + 10 bp terminal + 5 bp jitter. The SPRITE reads have some overrepresented nucleotides in cycles, so we added 20% of another library or PhiX to sequence. Quality control checks were performed on the final library pools using a Miseq nano kit according to the manufacturer’s instructions. Sequencing was performed using a Novaseq 6000 (Illumina) through Princeton’s Genomics Core Facility.

### RNA-SPRITE data pre-processing

A custom snakemake pipeline was used to process RNA-SPRITE sequencing data: https://github.com/SoftLivingMatter/RNA-SPRITE-Becker-2025. Briefly, data quality was assessed with fastqc (v0.12.1), barcodes were identified, and sequences without full barcodes were discarded (BarcodeIdentification_v1.2.0)^70^. MarkDuplicates (Picard) was used to remove PCR duplicates by filtering out similar sequences with the same barcode. Sequencing adapters and barcodes were removed before doing two pass RNAseq alignment. As described above, we performed paired-end sequencing with most of one read corresponding to the cluster barcode, therefore, alignments were based on approximately 150-200 nucleotides of sequencing. To avoid small, highly abundant RNAs spuriously aligning to mRNAs, we first aligned all sequences using Bowtie2 (v2.4.1) to a custom transcriptome with highly abundant non-coding RNAs (ncRNAs): snoRNAs, snRNAs, and rRNAs. We then took the unaligned reads from the first pass alignment and aligned them to a repeat-masked genome (GRCh38 or mRatBN7.2) using STAR (v2.7.11).

Further data processing and analysis was done using custom R scripts (https://github.com/SoftLivingMatter/RNA-SPRITE-Becker-2025). Briefly, Ensembl transcript ID alignments were aggregated together by their associated Ensembl gene ID, and all analyses were performed at the level gene ID. The Ensembl transcript ID sequence with the best coverage of alignments was chosen as the one representative sequence for each Ensembl gene ID. The gene IDs in each dataset were filtered to be above a minimum number of observations (unique barcodes) cutoff. We did not allow for homotypic colocalization; each barcode could only map to a gene once. We made a gene by cluster matrix only using clusters mapping to 2-20 genes of interest. For cytoplasmic mRNA analyses, we removed all ncRNAs, mRNAs encoded by mitochondrial DNA, and all mRNAs in clusters that also had alignments to known nuclear RNAs (snRNAs, snoRNAs, scaRNAs, pre-rRNA, and RMRP).

With our current technique, we can’t easily tell if two reads with the same barcode that align to different regions of the same transcript sequence are from a single RNA molecule or from two separate RNA copies from the same gene. RNA molecules are fragmented during the RNA-SPRITE protocol, and therefore a single RNA molecule can have multiple barcodes ligated to it in different locations along the transcript. To overcome this issue and a similar problem with homologs, we take a very conservative approach and ignore all colocalizations between transcripts that share sequence homology. As a result, we can’t report colocalization of multiple copies of a single RNA species. To avoid reporting colocalizations between transcripts with sequence homology, we aggregate observations aligned to paralogs into a single gene ID. We identified paralogs through Ensembl annotation (at least 40% sequence identity). Importantly, to ensure that none of the RNAs we found to be significantly colocalizing had strong sequence homology, we also checked all RNA pairs with a NBR raw z-score above 3 for sequence homology using Biostrings (v2.70.3) pairwiseAlignment function (“local-global” and “overlap”). For gene pairs with strong sequence homology (“local-global” > 0 or “overlap” > 50), their observations were aggregated into a single gene ID as well, and NBR of the dataset was repeated until no significantly colocalizing pairs demonstrated strong sequence homology.

### RNAseq and analysis

Whole neuron, total RNA isolation and library prep was performed as previously described^141^, by standard protocols. Alignment to the transcriptome was performed as described above in the RNA-SPRITE pipeline using a “two pass” alignment method. Ensembl transcript IDs were collapsed into Ensembl gene IDs, and gene IDs with high sequence homology were collapsed into a single gene ID as described above, in the same way as for RNA-SPRITE.

### mRNA colocalization propensity analysis

For each protein-coding gene, the number of colocalizations it had with another protein-coding gene was divided by the total number of observations. For example, if 3 different mRNAs aligned with the same barcode, this would be counted as 2 colocalizations for each mRNA. Observations include barcodes that only mapped to a single gene. Abundance, calculated as transcripts per million (TPM), was well correlated between RNA-SPRITE and RNAseq (Figure S1G, Spearman rho = 0.94). For each dataset, the mean colocalization propensity was set to 1 for ease of interpretation (by dividing every value by the mean value). For perturbation datasets and their paired, untreated experimental control datasets, the mean colocalization propensity of the control dataset was set to 1.

### RNA-binding protein (RBP) motif analysis

RBP motifs from ATtRACT^97^ and mCross^98^ databases were used if they were associated with an RBP whose encoding mRNA had an expression level above 5 TPM in the relevant (HEK or whole neuron) RNA-SPRITE dataset, to ensure only motifs for expressed RBPs were examined. The whole neuron 5 TPM cutoff was used for the neurite RNA-SPRITE dataset as well. Motif occurrences were called using FIMO (v4.11.2)^99^. SEA (v5.5.7) was used to calculate motif enrichment in a set of sequences^100^ vs. length-matched control sequences, 5% FDR (Benjamini Hochberg) cutoff. Putative liquid condensate enriched mRNAs were identified by being in the top quartile of mRNAs sensitive to hypotonic treatment (mRNA colocalization propensity in control - hypotonic conditions), and top quartile of mRNA colocalization propensity in control conditions (Figure S3D). For Figure S6G, an mRNA was considered a target of an RBP if it had two significant motifs for the RBP within 200 nt of each other in the whole transcript. All other analyses were done only using 3’ UTR sequences, and a single significant motif for an RBP was sufficient to be called as a target of an RBP. Only 3’ UTR sequences with length greater than 7 were analyzed.

### Dataset annotations

mRNA localization data were used from previous studies^40,58,60,67,76–79^. An mRNA was considered annotated as localized to a particular region if observed there in two assays, whenever at least two assays were available. Protein interaction data was used from CORUM, Complex Portal, and STRINGdb databases^111–113^. For STRINGdb, we used a physical score threshold of 700 as an annotation of a physical interaction. Proteins were considered interacting if they or their human homologs were identified as interacting in at least two databases. mRNA-mRNA duplexes were identified using the RISE database^107^. Only experiments that identified direct (not protein-dependent) RNA-RNA interactions were used: PARIS, LIGRseq, and SPLASH^63,64,107,108^. Interactions annotated between rodent or human protein-coding genes were used, but “intronic” or “intergenic” were not used. There was not enough overlap among the datasets to do a 2 assay cutoff as we did for other annotations.

### Free energy of folding estimates

Gibbs free energy of folding (ΔG) was estimated using NUPACK 4.0^88^ using custom scripts (https://github.com/SoftLivingMatter/RNA-SPRITE-Becker-2025). Free energy of dimerization was calculated by subtracting the summed monomer folding free energies from the dimer free energy. For free energy of dimerization comparisons, we considered the top 1000 highest confidence colocalizing mRNA pairs (by z-score) in each dataset vs. control pairs where the second mRNA was replaced with a length-matched control that did not colocalize with the first mRNA more than expected by chance. We also calculated the free energy of dimerization for colocalizing mRNA pairs found to form a duplex by proximity ligation experiments^107^ that also colocalized in HEK RNA-SPRITE data (850 pairs), and compared them to length-matched controls as above.

### Negative binomial regression

Motivated by its successful use in single-cell RNA-sequencing data analysis^75^, we applied negative binomial regression (NBR, formally a generalized linear model, GLM) to model pairwise counts. Given the scale of the data, we developed a custom R package for fitting (https://github.com/davidaknowles/adaglm). Briefly, adaglm fits a NBR using Adam, an algorithm for stochastic gradient descent optimization. Since Adam supports minibatching, adaglm can run on very large datasets (tens of millions of data points). By default the fitting is done in two steps: 1) Fit a Poisson regression. 2) Starting with coefficients initialized from the Poisson regression, fit a NBR jointly optimizing the coefficients and dispersion parameter. This two step procedure avoids inefficient optimization dynamics that can result from attempting to learn the regression coefficients and the dispersion parameter simultaneously.

NBR models the observed colocalization count *y*_*ab’*_ between RNAs *a* and *b*, as negative binomially distributed with expectation µ_*ab*_ and variance *var*(*y*_*ab*_) = µ_*ab*_ + γµ_*ab*_^2^ where γ_*ab*_ is the *dispersion* parameter which is shared across all pairs (note for γ = 0 the negative binomial is equivalent to a Poisson). In the baseline model, the expectation is determined by covariates *X*_*ab*_ according to,

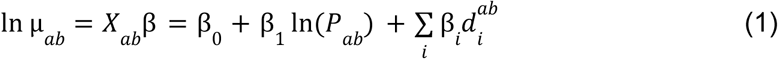

where β are learned regression coefficients and *P*_*ab*_ is the preliminarily expected colocalization count between the RNA pair calculated as *P*_*ab*_ = *y*_*a*•_ *y*_•*b*_ /*y*_••_ with *y*_*a*•_ and *y*_•*b*_ being the total colocalizations for RNA *a* and *b* respectively, and *y*_••_ being the total colocalization count in the dataset. *d*^ab^_*i*_ is a binary indicator denoting whether the genomic distance between *a* and *b* falls into distance bin *i.* The genomic distance bins refer to the distance between the genes encoding the two RNAs and whether they are encoded on the same DNA strand or the opposite strand. Eight genomic distance bins were used for NBR with the HEK and whole neuron datasets: 0 to 10 kb same strand, 10 to 50 kb same strand, 50 to 100 kb same strand, 100 to 500 kb same strand, 0 to 10 kb opposite strand, 10 to 50 kb opposite strand, 50 to 100 kb opposite strand, and 100 to 500 kb opposite strand. Only two genomic distance bins were used for the isolated neurite dataset because the other genomic bin terms were not significant in the NBR model: 0 to 10 kb same strand and 10 to 50 kb same strand.

Prior SPRITE studies looking at RNA-DNA colocalization demonstrated that co-transcriptional nucleic acid colocalizations can be picked up by SPRITE, and the increased colocalization for RNAs encoded within 500 kb (roughly exponentially decreasing with distance) are consistent with expectation for chromatin territories. Because the neurite dataset did not include material from cell nuclei, we believe that most of the increase in colocalizations between RNAs encoded within 50 kb on the same strand are due to the STAR alignment algorithm randomly choosing a transcript annotation when a sequencing read aligns equally well to multiple transcripts that are annotated for the same genomic locus. This is a common occurrence as many genomic loci, especially those that are highly transcriptionally active, can have many overlapping annotations with different Ensembl gene IDs. The genomic distance over which we observe this (0 to 50 kb) is consistent with the typical length of mammalian genes. As described above, we do not allow for homotypic colocalization (multiple alignments of a single gene ID to the same barcode) because we cannot discriminate whether the reads came from a single RNA molecule or separate molecules. However, that filter does not work if STAR calls one read as aligning to gene ID A and a second read from the same molecule as aligning to gene ID B. We used the genomic distance bins in the NBR to overcome this issue and account for the spurious increase in colocalizations between overlapping genes, as well as bona fide co-transcriptional colocalizations so that the genomic position of an RNA did not confound our analyses.

We defined colocalization “pairwise specificity” as the *Pearson residual* from the NBR,

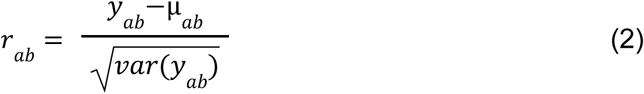

where µ_*ab*_ is from the model fit in Equation (1). *r*_*ab*_ reflects the degree to which the number of colocalizations between a given pair of RNAs deviates from an expectation of random colocalization. We calculated “within group” mean pairwise specificity by taking each RNA (mRNA or ncRNA, as labeled) annotated to localize to a particular subcellular region from other studies (see above) and calculating the mean pairwise specificity for that RNA with all other RNAs in the group (mRNAs and ncRNAs). “Outside group” mean pairwise specificity was calculated by taking each RNA (mRNA or ncRNA, as labeled) annotated to localize to a particular subcellular region and calculating the mean pairwise specificity for that RNA with all other RNAs *not* annotated to be in the group (mRNAs and ncRNAs). To assess the statistical significance of individual colocalization counts *y*_*ab*_, we calculated two-sided *p-*values taking the predictive distribution of Equation (1) as the null distribution. We convert these *p*-values into corresponding signed *z*-scores.

To estimate the effect of pairwise binary features *x*_*ab’*_, e.g., known protein-protein interaction (PPI) between the proteins encoded by *a* and *b*, we fit the extended NBR model,

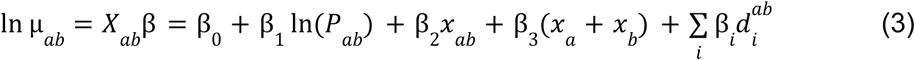

where *x_a_* and *x_b_* are binary indicators for *a* and *b* having an annotation in that category respectively (e.g., having any known PPI). The *x_a_* + *x_b_* term controls for any non-specific effect of the annotation. The remaining terms are analogous to those in the null model. We obtain standard errors for each coefficient β and use these to calculate Wald statistic *p*-values for the significance of each term. To ease interpretation, we calculate α = exp(β_2_) which is a multiplicative factor representing how much more frequent colocalization is when *x*_*ab*_ = 1. For the plots with α, the p-values reported with refer to the Wald statistic for β_2_, and the 95% confidence intervals = exp(β_2_ ± 2*standard error). We are primarily interested in the effect size reflected by α and whether it is statistically significantly different from 1, which implies the pairwise feature has a true effect on colocalization.

For testing whether shared binding of a specific RBP effects colocalization we use the extended NBR of Equation (3) where now *x*_*ab*_ = *x*_*a*_*x*_*b*_ = 1 if and only if the RBP binds both *a* and *b.* The control term *x*_*a*_ + *x*_*b*_ accounts for the possibility that binding of this RBP increases colocalization counts globally rather than specifically for partners to whom it also binds.

### mRNA colocalization hub analysis

mRNA-mRNA colocalization counts in the whole cell neuron dataset were analyzed using custom R scripts. Barcodes aligning to any known nuclear RNAs were excluded to focus on cytoplasmic organization of mature mRNAs. To examine only mRNAs that had evidence of specific colocalization, only mRNAs with a maximum pairwise colocalization z-score of at least 3.2 were used (7702 out of 8322 mRNAs). Negative pairwise specificity scores were changed to zero to avoid modeling deviation from expectation in non-colocalizing pairs.

To reduce noise and extract key axes of variation, we performed singular value decomposition (SVD) of the pairwise specificity scores using the RSpectra package (v0.16-2) in R, which efficiently computes the top K singular values and vectors for large matrices. The top K=5 left singular vectors (*U*) were scaled by their corresponding singular values (*D*) to obtain the SVD-transformed data. To identify local structure, we constructed a nearest neighbor graph using the RANN package (v2.6.2), connecting each mRNA to its 100 nearest neighbors. The resulting adjacency matrix, where entries were set to 1 for nearest neighbors, was converted into a graph object weighted by Pearson residuals. We then applied Louvain community detection (cluster_louvain function in igraph v2.1.4), using a resolution parameter of 0.5 to identify clusters.

Separately, we applied Uniform Manifold Approximation and Projection (UMAP) to the SVD-transformed data for nonlinear dimensionality reduction using the umap R package (v0.2.10.0). The embedding was generated using a cosine distance metric and 50 nearest neighbors, allowing for visualization of mRNA colocalization hubs.

### Gene ontology (GO) analysis

Custom R scripts with clusterProfiler^142^ (v3.20) were used for GO analysis. A 5% false discovery rate (FDR) cutoff was used and GO terms for biological process, molecular function, and cellular components were analyzed together for enrichment in each hub (Figure 3C). Hubs 4, 9, and 10 had the greatest number of significant GO terms. For these hubs, the simplify function with a 70% identity cutoff was used to eliminate redundant GO terms, keeping the GO term with lowest q-value. Then the top 10 GO terms by lowest q-value were used for plotting. For all other hubs, all significant GO terms with a 5% FDR are plotted. Hubs 1, 3, 6, and 8 had no significant GO terms.

### Deep learning models

We aimed to develop a deep learning (DL) model to predict colocalization specificity from the RNA sequences of the two colocalizing species. Custom scripts can be found at https://github.com/davidaknowles/locnet. DL models including convolutional neural networks^143–145^ (CNN), recurrent neural networks^146,147^ (RNN), transformers, and hybrids of these models^148^, have had substantial success as “sequence-to-function” models predicting diverse molecular phenotypes including transcription factor binding, chromatin accessibility, RBP binding and splicing. However, these existing models only take one sequence as input, rather than two as required in our case, and cannot easily handle varying length inputs. We therefore developed a novel neural network architecture. First, the network embeds both sequences into corresponding latent representations (K-dimensional vectors), using the same subnetwork for each. After exploring multiple choices for this subnetwork, including RNNs, we settled on using several convolutional layers with max-pooling followed by a single multi-headed attention (MHA) layer to summarize across the sequence dimension and produce the 32d vector. Second, the network calculates a weighted inner product of the two vectors, which is used as the prediction for colocalization specificity. We considered alternatives including a bilinear form with a K×K symmetric weight matrix, but did not see improvements in predictive performance. To represent an mRNA sequence of length L, we construct a 10kb one-hot encoding (i.e., with four channels corresponding to the four nucleotides) where the 5’ end corresponds to the first L/2 nucleotides, and the 3’ end corresponds to the last L/2 nucleotides. For rare very long mRNAs where L>10kb, some sequence in the middle of the mRNA will not be used. We ensure that the model only attends to the parts of the 10kb input that are not missing (i.e., correspond to actual mRNA sequence) by using the masking mechanism of MHA. This representation allows the model to reason about the relevance of different motifs at varying distances from the 5’ or 3’ end of the mRNA, and additionally simplifies saliency analysis.

We train the model to predict asinh-transformed Pearson residuals from the NBR so that total colocalizations per mRNA and genomic distance are already accounted for. asinh transformation is used to reduce the influence of rare large residuals. asinh behaves like log for large absolute values, linearly around 0, and is antisymmetric around 0. We use backpropagation and the Adam optimizer^149^ in torch v2.3.1 to train the network parameters, including the inner product weights, minimizing mean-squared error loss. We split the data randomly into 80% training, 10% validation and 10% test. Validation data was just for early stopping and hyperparameter optimization. We used ray.tune v1.13.0 ^150^ to select optimal hyperparameters: learning rate of 1e-3, dropout rate of 0.1, 5 convolutional layers, 16 hidden channels, filter width of 7, max-pooling stride of 3, 4 heads in the MHA and latent dimension K=32.

To understand the sequence features that the model learns to use to predict colocalization, we used saliency analysis, an approach adapted from computer vision^118^. The first convolutional layer of the network acts by scanning along the input sequence scoring how well each position matches to the learned convolutional filters (i.e., sequence motifs represented as position weight matrices, PWMs). These scores are passed through a rectified linear unit (ReLU) which zeros out negative entries to produce the first layer activations. For every positively colocalizing pair A and B (Pearson residual > 0) in the dataset we used backpropagation to calculate the gradient of the model output with respect to the first layer activations. The “saliency map” is defined as the gradient multiplied by the activation, and represents how much each motif match, at each position, contributes to the colocalization prediction for mRNA pair A-B. We average these saliency maps across pairs per sequence position to produce a 10kb average saliency map for each motif. Due to our strategy of placing the 5’ and 3’ mRNA sequence at the start and end of the 10kb input, we can directly read off the motif contribution at different distances from the 5’ or 3’ end.

We matched learned PWMs from the first to our curated set of known RBP motifs (see RNA-binding protein motif analysis above) accounting for differences in length by 0-padding and allowing position offsets in the match of up to 3 nucleotides.

### High performance computing

The analyses presented in this article were performed on computational resources managed and supported by Princeton Research Computing, a consortium of groups including the Princeton Institute for Computational Science and Engineering (PICSciE).

### Declaration of generative AI and AI-assisted technologies in the writing process

During the preparation of this work the authors used ChatGPT in order to improve clarity and readability of individual sections. After using this tool/service, the authors reviewed and edited the content as needed and take full responsibility for the content of the publication.

## REFERENCES

1. Antifeeva, I.A., Fonin, A.V., Fefilova, A.S., Stepanenko, O.V., Povarova, O.I., Silonov, S.A., Kuznetsova, I.M., Uversky, V.N., and Turoverov, K.K. (2022). Liquid-liquid phase separation as an organizing principle of intracellular space: overview of the evolution of the cell compartmentalization concept. Cell. Mol. Life Sci. 79, 251.

2. Kang, J.-Y., Wen, Z., Pan, D., Zhang, Y., Li, Q., Zhong, A., Yu, X., Wu, Y.-C., Chen, Y., Zhang, X., et al. (2022). LLPS of FXR1 drives spermiogenesis by activating translation of stored mRNAs. Science 377, eabj6647.

3. Ma, W., and Mayr, C. (2018). A Membraneless Organelle Associated with the Endoplasmic Reticulum Enables 3’UTR-Mediated Protein-Protein Interactions. Cell 175, 1492–1506.e19.

4. Cioni, J.-M., Lin, J.Q., Holtermann, A.V., Koppers, M., Jakobs, M.A.H., Azizi, A., Turner-Bridger, B., Shigeoka, T., Franze, K., Harris, W.A., et al. (2019). Late endosomes act as mRNA translation platforms and sustain mitochondria in axons. Cell 176, 56–72.e15.

5. Forrest, K.M., and Gavis, E.R. (2003). Live imaging of endogenous RNA reveals a diffusion and entrapment mechanism for nanos mRNA localization in Drosophila. Curr. Biol. 13, 1159–1168.

6. Bose, M., Lampe, M., Mahamid, J., and Ephrussi, A. (2022). Liquid-to-solid phase transition of oskar ribonucleoprotein granules is essential for their function in Drosophila embryonic development. Cell 185, 1308–1324.e23.

7. Lee, C.-Y.S., Putnam, A., Lu, T., He, S., Ouyang, J.P.T., and Seydoux, G. (2020). Recruitment of mRNAs to P granules by condensation with intrinsically-disordered proteins. Elife 9. 10.7554/eLife.52896.

8. Di Stefano, B., Luo, E.-C., Haggerty, C., Aigner, S., Charlton, J., Brumbaugh, J., Ji, F., Rabano Jiménez, I., Clowers, K.J., Huebner, A.J., et al. (2019). The RNA helicase DDX6 controls cellular plasticity by modulating P-body homeostasis. Cell Stem Cell 25, 622–638.e13.

9. Liao, Y.-C., Fernandopulle, M.S., Wang, G., Choi, H., Hao, L., Drerup, C.M., Patel, R., Qamar, S., Nixon-Abell, J., Shen, Y., et al. (2019). RNA granules hitchhike on lysosomes for long-distance transport, using annexin A11 as a molecular tether. Cell 179, 147–164.e20.

10. Cardona, A.H., Ecsedi, S., Khier, M., Yi, Z., Bahri, A., Ouertani, A., Valero, F., Labrosse, M., Rouquet, S., Robert, S., et al. (2023). Self-demixing of mRNA copies buffers mRNA:mRNA and mRNA:regulator stoichiometries. Cell 186, 4310–4324.e23.

11. Boke, E., Ruer, M., Wühr, M., Coughlin, M., Lemaitre, R., Gygi, S.P., Alberti, S., Drechsel, D., Hyman, A.A., and Mitchison, T.J. (2016). Amyloid-like self-assembly of a cellular compartment. Cell 166, 637–650.

12. An, J.J., Gharami, K., Liao, G.-Y., Woo, N.H., Lau, A.G., Vanevski, F., Torre, E.R., Jones, K.R., Feng, Y., Lu, B., et al. (2008). Distinct role of long 3’ UTR BDNF mRNA in spine morphology and synaptic plasticity in hippocampal neurons. Cell 134, 175–187.

13. Miller, S., Yasuda, M., Coats, J.K., Jones, Y., Martone, M.E., and Mayford, M. (2002). Disruption of dendritic translation of CaMKIIα impairs stabilization of synaptic plasticity and memory consolidation. Neuron 36, 507–519.

14. Bourke, A.M., Schwarz, A., and Schuman, E.M. (2023). De-centralizing the Central Dogma: mRNA translation in space and time. Mol Cell 83, 452–468.

15. Shapiro, D.M., Deshpande, S., Eghtesadi, S.A., Zhong, M., Fontes, C.M., Fiflis, D., Rohm, D., Min, J., Kaur, T., Peng, J., et al. (2025). Synthetic biomolecular condensates enhance translation from a target mRNA in living cells. Nat. Chem. 10.1038/s41557-024-01706-7.

16. Roden, C., and Gladfelter, A.S. (2021). RNA contributions to the form and function of biomolecular condensates. Nat Rev Mol Cell Biol 22, 183–195.

17. Shin, Y., and Brangwynne, C.P. (2017). Liquid phase condensation in cell physiology and disease. Science. 10.1126/science.aaf4382.

18. Jain, A., and Vale, R.D. (2017). RNA phase transitions in repeat expansion disorders. Nature 546, 243–247.

19. Zhang, H., Elbaum-Garfinkle, S., Langdon, E.M., Taylor, N., Occhipinti, P., Bridges, A.A., Brangwynne, C.P., and Gladfelter, A.S. (2015). RNA controls PolyQ protein phase transitions. Mol. Cell 60, 220–230.

20. Elbaum-Garfinkle, S., Kim, Y., Szczepaniak, K., Chen, C.C.-H., Eckmann, C.R., Myong, S., and Brangwynne, C.P. (2015). The disordered P granule protein LAF-1 drives phase separation into droplets with tunable viscosity and dynamics. Proc. Natl. Acad. Sci. U. S. A. 112, 7189–7194.

21. Boeynaems, S., Holehouse, A.S., Weinhardt, V., Kovacs, D., Van Lindt, J., Larabell, C., Van Den Bosch, L., Das, R., Tompa, P.S., Pappu, R.V., et al. (2019). Spontaneous driving forces give rise to protein-RNA condensates with coexisting phases and complex material properties. Proc. Natl. Acad. Sci. U. S. A. 116, 7889–7898.

22. Mann, J.R., Gleixner, A.M., Mauna, J.C., Gomes, E., DeChellis-Marks, M.R., Needham, P.G., Copley, K.E., Hurtle, B., Portz, B., Pyles, N.J., et al. (2019). RNA binding antagonizes neurotoxic phase transitions of TDP-43. Neuron 102, 321–338.e8.

23. Van Treeck, B., Protter, D.S.W., Matheny, T., Khong, A., Link, C.D., and Parker, R. (2018). RNA self-assembly contributes to stress granule formation and defining the stress granule transcriptome. Proc. Natl. Acad. Sci. U. S. A. 115, 2734–2739.

24. Ambadi Thody, S., Clements, H.D., Baniasadi, H., Lyon, A.S., Sigman, M.S., and Rosen, M.K. (2024). Small-molecule properties define partitioning into biomolecular condensates. Nat Chem 16, 1794–1802.

25. Matthew R King, Kiersten M Ruff, Andrew Z Lin, Avnika Pant, Mina Farag, Jared M Lalmansingh, Tingting Wu, Martin J Fossat, Wei Ouyang, Matthew D Lew, Emma Lundberg, Michael D Vahey, Rohit V Pappu (2024). Macromolecular condensation organizes nucleolar sub-phases to set up a pH gradient. Cell 187, 1889–1906.e24.

26. Ausserwöger, H., Scrutton, R., Sneideris, T., Fischer, C.M., Qian, D., de Csilléry, E., Saar, K.L., Białek, A.Z., Oeller, M., Krainer, G., et al. (2024). Biomolecular condensates sustain pH gradients at equilibrium driven by charge neutralisation. bioRxiv, 2024.05.23.595321. 10.1101/2024.05.23.595321.

27. Nott, T.J., Craggs, T.D., and Baldwin, A.J. (2016). Membraneless organelles can melt nucleic acid duplexes and act as biomolecular filters. Nat. Chem. 8, 569–575.

28. Choi, S., Meyer, M.O., Bevilacqua, P.C., and Keating, C.D. (2022). Phase-specific RNA accumulation and duplex thermodynamics in multiphase coacervate models for membraneless organelles. Nat. Chem. 14, 1110–1117.

29. Kim, T.H., Tsang, B., Vernon, R.M., Sonenberg, N., Kay, L.E., and Forman-Kay, J.D. (2019). Phospho-dependent phase separation of FMRP and CAPRIN1 recapitulates regulation of translation and deadenylation. Science 365, 825–829.

30. Tibble, R.W., Depaix, A., Kowalska, J., Jemielity, J., and Gross, J.D. (2021). Biomolecular condensates amplify mRNA decapping by biasing enzyme conformation. Nat. Chem. Biol. 17, 615–623.

31. Shen, C., Li, R., Negro, R., Cheng, J., Vora, S.M., Fu, T.-M., Wang, A., He, K., Andreeva, L., Gao, P., et al. (2021). Phase separation drives RNA virus-induced activation of the NLRP6 inflammasome. Cell 184, 5759–5774.e20.

32. Strzelecka, M., Trowitzsch, S., Weber, G., Lührmann, R., Oates, A.C., and Neugebauer, K.M. (2010). Coilin-dependent snRNP assembly is essential for zebrafish embryogenesis. Nat. Struct. Mol. Biol. 17, 403–409.

33. Holt, C.E., Martin, K.C., and Schuman, E.M. (2019). Local translation in neurons: visualization and function. Nat Struct Mol Biol 26, 557–566.

34. Costa-Mattioli, M., Sossin, W.S., Klann, E., and Sonenberg, N. (2009). Translational control of long-lasting synaptic plasticity and memory. Neuron 61, 10–26.

35. Rangaraju, V., Lauterbach, M., and Schuman, E.M. (2019). Spatially stable mitochondrial compartments fuel local translation during plasticity. Cell 176, 73–84.e15.

36. Bauer, K.E., de Queiroz, B.R., Kiebler, M.A., and Besse, F. (2023). RNA granules in neuronal plasticity and disease. Trends Neurosci. 46, 525–538.

37. Kiebler, M.A., and Bauer, K.E. (2024). RNA granules in flux: dynamics to balance physiology and pathology. Nat. Rev. Neurosci. 25, 711–725.

38. Monday, H.R., Kharod, S.C., Yoon, Y.J., Singer, R.H., and Castillo, P.E. (2022). Presynaptic FMRP and local protein synthesis support structural and functional plasticity of glutamatergic axon terminals. Neuron 110, 2588–2606.e6.

39. Hafner, A.-S., Donlin-Asp, P.G., Leitch, B., Herzog, E., and Schuman, E.M. (2019). Local protein synthesis is a ubiquitous feature of neuronal pre- and postsynaptic compartments. Science 364. 10.1126/science.aau3644.

40. Chen, X., Fansler, M.M., Janjoš, U., Ule, J., and Mayr, C. (2024). The FXR1 network acts as a signaling scaffold for actomyosin remodeling. Cell 187, 5048–5063.e25.

41. Yan, X., Kuster, D., Mohanty, P., Nijssen, J., Pombo-García, K., Rizuan, A., Franzmann, T.M., Sergeeva, A., Passos, P.M., George, L., et al. (2024). Intra-condensate demixing of TDP-43 inside stress granules generates pathological aggregates. bioRxiv. 10.1101/2024.01.23.576837.

42. Zhang, P., Fan, B., Yang, P., Temirov, J., Messing, J., Kim, H.J., and Taylor, J.P. (2019). Chronic optogenetic induction of stress granules is cytotoxic and reveals the evolution of ALS-FTD pathology. Elife 8. 10.7554/eLife.39578.

43. Molliex, A., Temirov, J., Lee, J., Coughlin, M., Kanagaraj, A.P., Kim, H.J., Mittag, T., and Taylor, J.P. (2015). Phase separation by low complexity domains promotes stress granule assembly and drives pathological fibrillization. Cell 163, 123–133.

44. Patel, A., Lee, H.O., Jawerth, L., Maharana, S., Jahnel, M., Hein, M.Y., Stoynov, S., Mahamid, J., Saha, S., Franzmann, T.M., et al. (2015). A liquid-to-solid phase transition of the ALS protein FUS accelerated by disease mutation. Cell 162, 1066–1077.

45. Lee, K.-H., Zhang, P., Kim, H.J., Mitrea, D.M., Sarkar, M., Freibaum, B.D., Cika, J., Coughlin, M., Messing, J., Molliex, A., et al. (2016). C9orf72 dipeptide repeats impair the assembly, dynamics, and function of membrane-less organelles. Cell 167, 774–788.e17.

46. Alami, N.H., Smith, R.B., Carrasco, M.A., Williams, L.A., Winborn, C.S., Han, S.S.W., Kiskinis, E., Winborn, B., Freibaum, B.D., Kanagaraj, A., et al. (2014). Axonal transport of TDP-43 mRNA granules is impaired by ALS-causing mutations. Neuron 81, 536–543.

47. Nagano, S., Jinno, J., Abdelhamid, R.F., Jin, Y., Shibata, M., Watanabe, S., Hirokawa, S., Nishizawa, M., Sakimura, K., Onodera, O., et al. (2020). TDP-43 transports ribosomal protein mRNA to regulate axonal local translation in neuronal axons. Acta Neuropathol. 140, 695–713.

48. Chu, J.-F., Majumder, P., Chatterjee, B., Huang, S.-L., and Shen, C.-K.J. (2019). TDP-43 regulates coupled dendritic mRNA transport-translation processes in co-operation with FMRP and Staufen1. Cell Rep. 29, 3118–3133.e6.

49. Jinno, J., Abdelhamid, R.F., Morita, J., Saga, R., Yamasaki, Y., Kadowaki, A., Ogawa, K., Kimura, Y., Ikenaka, K., Beck, G., et al. (2025). TDP-43 transports ferritin heavy chain mRNA to regulate oxidative stress in neuronal axons. Neurochem. Int., 105934.

50. Combs, B., Christensen, K.R., Richards, C., Kneynsberg, A., Mueller, R.L., Morris, S.L., Morfini, G.A., Brady, S.T., and Kanaan, N.M. (2021). Frontotemporal lobar dementia mutant tau impairs axonal transport through a protein phosphatase 1γ-dependent mechanism. J. Neurosci. 41, 9431–9451.

51. Stokin, G.B., Lillo, C., Falzone, T.L., Brusch, R.G., Rockenstein, E., Mount, S.L., Raman, R., Davies, P., Masliah, E., Williams, D.S., et al. (2005). Axonopathy and transport deficits early in the pathogenesis of Alzheimer’s disease. Science 307, 1282–1288.

52. López-Erauskin, J., Tadokoro, T., Baughn, M.W., Myers, B., McAlonis-Downes, M., Chillon-Marinas, C., Asiaban, J.N., Artates, J., Bui, A.T., Vetto, A.P., et al. (2020). ALS/FTD-linked mutation in FUS suppresses intra-axonal protein synthesis and drives disease without nuclear loss-of-function of FUS. Neuron 106, 354.

53. Bakthavachalu, B., Huelsmeier, J., Sudhakaran, I.P., Hillebrand, J., Singh, A., Petrauskas, A., Thiagarajan, D., Sankaranarayanan, M., Mizoue, L., Anderson, E.N., et al. (2018). RNP-granule assembly via Ataxin-2 disordered domains is required for long-term memory and neurodegeneration. Neuron 98, 754–766.e4.

54. Wijegunawardana, D., Nayak, A., Vishal, S.S., Venkatesh, N., and Gopal, P.P. (2025). Ataxin-2 polyglutamine expansions aberrantly sequester TDP-43 ribonucleoprotein condensates disrupting mRNA transport and local translation in neurons. Dev. Cell 60, 253–269.e5.

55. Fukuda, Y., Pazyra-Murphy, M.F., Silagi, E.S., Tasdemir-Yilmaz, O.E., Li, Y., Rose, L., Yeoh, Z.C., Vangos, N.E., Geffken, E.A., Seo, H.-S., et al. (2021). Binding and transport of SFPQ-RNA granules by KIF5A/KLC1 motors promotes axon survival. J. Cell Biol. 220. 10.1083/jcb.202005051.

56. Guerra San Juan, I., Brunner, J.W., Eggan, K., Toonen, R.F., and Verhage, M. (2025). KIF5A regulates axonal repair and time-dependent axonal transport of SFPQ granules and mitochondria in human motor neurons. Neurobiol. Dis. 204, 106759.

57. Jung, Y., Seo, J.-Y., Ryu, H.G., Kim, D.-Y., Lee, K.-H., and Kim, K.-T. (2020). BDNF-induced local translation of GluA1 is regulated by HNRNP A2/B1. Sci. Adv. 6, eabd2163.

58. Fazal, F.M., Han, S., Parker, K.R., Kaewsapsak, P., Xu, J., Boettiger, A.N., Chang, H.Y., and Ting, A.Y. (2019). Atlas of subcellular RNA localization revealed by APEX-seq. Cell 178, 473–490.e26.

59. Licatalosi, D.D., Mele, A., Fak, J.J., Ule, J., Kayikci, M., Chi, S.W., Clark, T.A., Schweitzer, A.C., Blume, J.E., Wang, X., et al. (2008). HITS-CLIP yields genome-wide insights into brain alternative RNA processing. Nature 456, 464–469.

60. Hubstenberger, A., Courel, M., Bénard, M., Souquere, S., Ernoult-Lange, M., Chouaib, R., Yi, Z., Morlot, J.-B., Munier, A., Fradet, M., et al. (2017). P-body purification reveals the condensation of repressed mRNA regulons. Mol. Cell 68, 144–157.e5.

61. Cao, C., Cai, Z., Ye, R., Su, R., Hu, N., Zhao, H., and Xue, Y. (2021). Global in situ profiling of RNA-RNA spatial interactions with RIC-seq. Nat. Protoc. 16, 2916–2946.

62. Wu, T., Cheng, A.Y., Zhang, Y., Xu, J., Wu, J., Wen, L., Li, X., Liu, B., Dou, X., Wang, P., et al. (2024). KARR-seq reveals cellular higher-order RNA structures and RNA-RNA interactions. Nat. Biotechnol. 10.1038/s41587-023-02109-8.

63. Lu, Z., Zhang, Q.C., Lee, B., Flynn, R.A., Smith, M.A., Robinson, J.T., Davidovich, C., Gooding, A.R., Goodrich, K.J., Mattick, J.S., et al. (2016). RNA Duplex Map in Living Cells Reveals Higher-Order Transcriptome Structure. Cell 165, 1267–1279.

64. Sharma, E., Sterne-Weiler, T., O’Hanlon, D., and Blencowe, B.J. (2016). Global mapping of human RNA-RNA interactions. Mol. Cell 62, 618–626.

65. Nguyen, T.C., Cao, X., Yu, P., Xiao, S., Lu, J., Biase, F.H., Sridhar, B., Huang, N., Zhang, K., and Zhong, S. (2016). Mapping RNA–RNA interactome and RNA structure in vivo by MARIO. Nat. Commun. 7, 12023.

66. Eng, C.-H.L., Lawson, M., Zhu, Q., Dries, R., Koulena, N., Takei, Y., Yun, J., Cronin, C., Karp, C., Yuan, G.-C., et al. (2019). Transcriptome-scale super-resolved imaging in tissues by RNA seqFISH. Nature 568, 235–239.

67. Xia, C., Fan, J., Emanuel, G., Hao, J., and Zhuang, X. (2019). Spatial transcriptome profiling by MERFISH reveals subcellular RNA compartmentalization and cell cycle-dependent gene expression. Proc. Natl. Acad. Sci. U. S. A. 116, 19490–19499.

68. Bressan, D., Battistoni, G., and Hannon, G.J. (2023). The dawn of spatial omics. Science 381, eabq4964.

69. Wang, G., Moffitt, J.R., and Zhuang, X. (2018). Multiplexed imaging of high-density libraries of RNAs with MERFISH and expansion microscopy. Sci. Rep. 8, 4847.

70. Quinodoz, S.A., Ollikainen, N., Tabak, B., Palla, A., Schmidt, J.M., Detmar, E., Lai, M.M., Shishkin, A.A., Bhat, P., Takei, Y., et al. (2018). Higher-Order Inter-chromosomal Hubs Shape 3D Genome Organization in the Nucleus. Cell 174, 744–757.e24.

71. Quinodoz, S.A., Jachowicz, J.W., Bhat, P., Ollikainen, N., Banerjee, A.K., Goronzy, I.N., Blanco, M.R., Chovanec, P., Chow, A., Markaki, Y., et al. (2021). RNA promotes the formation of spatial compartments in the nucleus. Cell 184, 5775–5790.e30.

72. Goronzy, I.N., Quinodoz, S.A., Jachowicz, J.W., Ollikainen, N., Bhat, P., and Guttman, M. (2022). Simultaneous mapping of 3D structure and nascent RNAs argues against nuclear compartments that preclude transcription. Cell Rep. 41, 111730.

73. Quinodoz, S.A., Bhat, P., Chovanec, P., Jachowicz, J.W., Ollikainen, N., Detmar, E., Soehalim, E., and Guttman, M. (2022). SPRITE: a genome-wide method for mapping higher-order 3D interactions in the nucleus using combinatorial split-and-pool barcoding. Nat. Protoc. 17, 36–75.

74. Khong, A., Matheny, T., Jain, S., Mitchell, S.F., Wheeler, J.R., and Parker, R. (2017). The stress granule transcriptome reveals principles of mRNA accumulation in stress granules. Mol. Cell 68, 808–820.e5.

75. Hafemeister, C., and Satija, R. (2019). Normalization and variance stabilization of single-cell RNA-seq data using regularized negative binomial regression. Genome Biol. 20, 296.

76. Horste, E.L., Fansler, M.M., Cai, T., Chen, X., Mitschka, S., Zhen, G., Lee, F.C.Y., Ule, J., and Mayr, C. (2023). Subcytoplasmic location of translation controls protein output. Mol. Cell 83, 4509–4523.e11.

77. Villanueva, E., Smith, T., Pizzinga, M., Elzek, M., Queiroz, R.M.L., Harvey, R.F., Breckels, L.M., Crook, O.M., Monti, M., Dezi, V., et al. (2024). System-wide analysis of RNA and protein subcellular localization dynamics. Nat. Methods 21, 60–71.

78. Barutcu, A.R., Wu, M., Braunschweig, U., Dyakov, B.J.A., Luo, Z., Turner, K.M., Durbic, T., Lin, Z.-Y., Weatheritt, R.J., Maass, P.G., et al. (2022). Systematic mapping of nuclear domain-associated transcripts reveals speckles and lamina as hubs of functionally distinct retained introns. Mol. Cell 82, 1035–1052.e9.

79. Ascano, M., Jr, Mukherjee, N., Bandaru, P., Miller, J.B., Nusbaum, J.D., Corcoran, D.L., Langlois, C., Munschauer, M., Dewell, S., Hafner, M., et al. (2012). FMRP targets distinct mRNA sequence elements to regulate protein expression. Nature 492, 382–386.

80. Padrón, A., Iwasaki, S., and Ingolia, N.T. (2019). Proximity RNA labeling by APEX-seq reveals the organization of translation initiation complexes and repressive RNA granules. Mol. Cell 75, 875–887.e5.

81. Ries, R.J., Pickering, B.F., Poh, H.X., Namkoong, S., and Jaffrey, S.R. (2023). m6A governs length-dependent enrichment of mRNAs in stress granules. Nat. Struct. Mol. Biol. 30, 1525–1535.

82. Glauninger, H., Bard, J.A.M., Wong Hickernell, C.J., Airoldi, E.M., Li, W., Singer, R.H., Paul, S., Fei, J., Sosnick, T.R., Wallace, E.W.J., et al. (2024). Transcriptome-wide mRNA condensation precedes stress granule formation and excludes stress-induced transcripts. bioRxiv. 10.1101/2024.04.15.589678.

83. Nott, T.J., Petsalaki, E., Farber, P., Jervis, D., Fussner, E., Plochowietz, A., Craggs, T.D., Bazett-Jones, D.P., Pawson, T., Forman-Kay, J.D., et al. (2015). Phase transition of a disordered nuage protein generates environmentally responsive membraneless organelles. Mol. Cell 57, 936–947.

84. Bounedjah, O., Hamon, L., Savarin, P., Desforges, B., Curmi, P.A., and Pastré, D. (2012). Macromolecular crowding regulates assembly of mRNA stress granules after osmotic stress: new role for compatible osmolytes. J. Biol. Chem. 287, 2446–2458.

85. Sehgal, P.B., Yuan, H., and Jin, Y. (2022). Rapid reversible osmoregulation of cytoplasmic biomolecular condensates of human interferon-α-induced antiviral MxA GTPase. Int. J. Mol. Sci. 23, 12739.

86. Blondel, V., Guillaume, J.-L., and Lambiotte, R. (2024). Fast unfolding of communities in large networks: 15 years later. J. Stat. Mech. 2024, 10R001.

87. McInnes, L., Healy, J., and Melville, J. (2018). UMAP: Uniform Manifold Approximation and Projection for Dimension Reduction. arXiv [stat.ML].

88. Fornace, M.E., Huang, J., Newman, C.T., Porubsky, N.J., Pierce, M.B., and Pierce, N.A. (2022). NUPACK: Analysis and design of nucleic acid structures, devices, and systems. ChemRxiv. 10.26434/chemrxiv-2022-xv98l.

89. Ma, W., Zhen, G., Xie, W., and Mayr, C. (2021). In vivo reconstitution finds multivalent RNA-RNA interactions as drivers of mesh-like condensates. Elife 10. 10.7554/eLife.64252.

90. Li, P., Banjade, S., Cheng, H.-C., Kim, S., Chen, B., Guo, L., Llaguno, M., Hollingsworth, J.V., King, D.S., Banani, S.F., et al. (2012). Phase transitions in the assembly of multivalent signalling proteins. Nature 483, 336–340.

91. Sanders, D.W., Kedersha, N., Lee, D.S.W., Strom, A.R., Drake, V., Riback, J.A., Bracha, D., Eeftens, J.M., Iwanicki, A., Wang, A., et al. (2020). Competing protein-RNA interaction networks control multiphase intracellular organization. Cell 181, 306–324.e28.

92. Boeynaems, S., Alberti, S., Fawzi, N.L., Mittag, T., Polymenidou, M., Rousseau, F., Schymkowitz, J., Shorter, J., Wolozin, B., Van Den Bosch, L., et al. (2018). Protein phase separation: A new phase in cell biology. Trends Cell Biol. 28, 420–435.

93. Dominguez, D., Freese, P., Alexis, M.S., Su, A., Hochman, M., Palden, T., Bazile, C., Lambert, N.J., Van Nostrand, E.L., Pratt, G.A., et al. (2018). Sequence, structure, and context preferences of human RNA binding proteins. Mol. Cell 70, 854–867.e9.

94. Li, X., Quon, G., Lipshitz, H.D., and Morris, Q. (2010). Predicting in vivo binding sites of RNA-binding proteins using mRNA secondary structure. RNA 16, 1096–1107.

95. Tushev, G., Glock, C., Heumüller, M., Biever, A., Jovanovic, M., and Schuman, E.M. (2018). Alternative 3′ UTRs modify the localization, regulatory potential, stability, and plasticity of mRNAs in neuronal compartments. Neuron 98, 495–511.e6.

96. Baltz, A.G., Munschauer, M., Schwanhäusser, B., Vasile, A., Murakawa, Y., Schueler, M., Youngs, N., Penfold-Brown, D., Drew, K., Milek, M., et al. (2012). The mRNA-bound proteome and its global occupancy profile on protein-coding transcripts. Mol. Cell 46, 674–690.

97. Giudice, G., Sánchez-Cabo, F., Torroja, C., and Lara-Pezzi, E. (2016). ATtRACT-a database of RNA-binding proteins and associated motifs. Database (Oxford) 2016, baw035.

98. Feng, H., Bao, S., Rahman, M.A., Weyn-Vanhentenryck, S.M., Khan, A., Wong, J., Shah, A., Flynn, E.D., Krainer, A.R., and Zhang, C. (2019). Modeling RNA-binding protein specificity in vivo by precisely registering protein-RNA crosslink sites. Mol. Cell 74, 1189–1204.e6.

99. Grant, C.E., Bailey, T.L., and Noble, W.S. (2011). FIMO: scanning for occurrences of a given motif. Bioinformatics 27, 1017–1018.

100. Bailey, T.L., and Grant, C.E. (2021). SEA: Simple Enrichment Analysis of motifs. bioRxiv. 10.1101/2021.08.23.457422.

101. Parker, D.M., Winkenbach, L.P., and Osborne Nishimura, E. (2022). It’s just a phase: Exploring the relationship between mRNA, biomolecular condensates, and translational control. Front. Genet. 13, 931220.

102. Venkataramanan, S., Gadek, M., Calviello, L., Wilkins, K., and Floor, S.N. (2021). DDX3X and DDX3Y are redundant in protein synthesis. RNA 27, 1577–1588.

103. Glock, C., Biever, A., Tushev, G., Nassim-Assir, B., Kao, A., Bartnik, I., Tom Dieck, S., and Schuman, E.M. (2021). The translatome of neuronal cell bodies, dendrites, and axons. Proc. Natl. Acad. Sci. U. S. A. 118, e2113929118.

104. Gao, X., Wan, J., Liu, B., Ma, M., Shen, B., and Qian, S.-B. (2015). Quantitative profiling of initiating ribosomes in vivo. Nat. Methods 12, 147–153.

105. Marklund, E., Ke, Y., and Greenleaf, W.J. (2023). High-throughput biochemistry in RNA sequence space: predicting structure and function. Nat. Rev. Genet. 24, 401–414.

106. Chouaib, R., Safieddine, A., Pichon, X., Imbert, A., Kwon, O.S., Samacoits, A., Traboulsi, A.-M., Robert, M.-C., Tsanov, N., Coleno, E., et al. (2020). A Dual Protein-mRNA Localization Screen Reveals Compartmentalized Translation and Widespread Co-translational RNA Targeting. Dev. Cell 54, 773–791.e5.

107. Gong, J., Shao, D., Xu, K., Lu, Z., Lu, Z.J., Yang, Y.T., and Zhang, Q.C. (2018). RISE: a database of RNA interactome from sequencing experiments. Nucleic Acids Res. 46, D194–D201.

108. Aw, J.G.A., Shen, Y., Wilm, A., Sun, M., Lim, X.N., Boon, K.-L., Tapsin, S., Chan, Y.-S., Tan, C.-P., Sim, A.Y.L., et al. (2016). In vivo mapping of eukaryotic RNA interactomes reveals principles of higher-order organization and regulation. Mol. Cell 62, 603–617.

109. Parker, D.M., Tauber, D., and Parker, R. (2025). G3BP1 promotes intermolecular RNA-RNA interactions during RNA condensation. Mol Cell. 10.1016/j.molcel.2024.11.012.

110. Trussina, I.R.E.A., Hartmann, A., Desroches Altamirano, C., Natarajan, J., Fischer, C.M., Aleksejczuk, M., Ausserwöger, H., Knowles, T.P.J., Schlierf, M., Franzmann, T.M., et al. (2025). G3BP-driven RNP granules promote inhibitory RNA-RNA interactions resolved by DDX3X to regulate mRNA translatability. Mol. Cell. 10.1016/j.molcel.2024.11.039.

111. Szklarczyk, D., Kirsch, R., Koutrouli, M., Nastou, K., Mehryary, F., Hachilif, R., Gable, A.L., Fang, T., Doncheva, N.T., Pyysalo, S., et al. (2023). The STRING database in 2023: protein-protein association networks and functional enrichment analyses for any sequenced genome of interest. Nucleic Acids Res. 51, D638–D646.

112. Meldal, B.H.M., Bye-A-Jee, H., Gajdoš, L., Hammerová, Z., Horácková, A., Melicher, F., Perfetto, L., Pokorný, D., Lopez, M.R., Türková, A., et al. (2019). Complex Portal 2018: extended content and enhanced visualization tools for macromolecular complexes. Nucleic Acids Res. 47, D550–D558.

113. Tsitsiridis, G., Steinkamp, R., Giurgiu, M., Brauner, B., Fobo, G., Frishman, G., Montrone, C., and Ruepp, A. (2023). CORUM: the comprehensive resource of mammalian protein complexes-2022. Nucleic Acids Res. 51, D539–D545.

114. Sahoo, P.K., Agrawal, M., Hanovice, N., Ward, P.J., Desai, M., Smith, T.P., SiMa, H., Dulin, J.N., Vaughn, L.S., Tuszynski, M.H., et al. (2025). Disruption of G3BP1 granules promotes mammalian CNS and PNS axon regeneration. Proc. Natl. Acad. Sci. U. S. A. 122, e2411811122.

115. Smith, R., Rathod, R.J., Rajkumar, S., and Kennedy, D. (2014). Nervous translation, do you get the message? A review of mRNPs, mRNA-protein interactions and translational control within cells of the nervous system. Cell. Mol. Life Sci. 71, 3917–3937.

116. Tarannum, R., Mun, G., Quddos, F., Swanger, S.A., Steward, O., and Farris, S. (2024). Dendritically localized RNAs are packaged as diversely composed ribonucleoprotein particles with heterogeneous copy number states. bioRxiv. 10.1101/2024.07.13.603387.

117. Markmiller, S., Soltanieh, S., Server, K.L., Mak, R., Jin, W., Fang, M.Y., Luo, E.-C., Krach, F., Yang, D., Sen, A., et al. (2018). Context-dependent and disease-specific diversity in protein interactions within stress granules. Cell 172, 590–604.e13.

118. Simonyan, K., Vedaldi, A., and Zisserman, A. (2013). Deep inside Convolutional Networks: Visualising image classification models and saliency maps. arXiv [cs.CV].

119. Shrikumar, A., Greenside, P., and Kundaje, A. (2017). Learning Important Features Through Propagating Activation Differences. In Proceedings of Machine Learning Research., D. Precup and Y. W. Teh, eds. (PMLR), pp. 3145–3153.

120. Chen, H., Lundberg, S.M., and Lee, S.-I. (2022). Explaining a series of models by propagating Shapley values. Nat. Commun. 13, 4512.

121. Ripin, N., and Parker, R. (2023). Formation, function, and pathology of RNP granules. Cell 186, 4737–4756.

122. Putnam, A., Thomas, L., and Seydoux, G. (2023). RNA granules: functional compartments or incidental condensates? Genes Dev. 37, 354–376.

123. de Vries, T., Novakovic, M., Ni, Y., Smok, I., Inghelram, C., Bikaki, M., Sarnowski, C.P., Han, Y., Emmanouilidis, L., Padroni, G., et al. (2024). Specific protein-RNA interactions are mostly preserved in biomolecular condensates. Sci. Adv. 10, eadm7435.

124. Quinodoz, S.A., Jiang, L., Abu-Alfa, A.A., Comi, T.J., Zhao, H., Yu, Q., Wiesner, L.W., Botello, J.F., Donlic, A., Soehalim, E., et al. (2024). Mapping and engineering RNA-controlled architecture of the multiphase nucleolus. bioRxivorg. 10.1101/2024.09.28.615444.

125. Wadsworth, G.M., Srinivasan, S., Lai, L.B., Datta, M., Gopalan, V., and Banerjee, P.R. (2024). RNA-driven phase transitions in biomolecular condensates. Mol. Cell 84, 3692–3705.

126. Seim, I., Zhang, V., Jalihal, A.P., Stormo, B.M., Cole, S.J., Ekena, J., Nguyen, H.T., Thirumalai, D., and Gladfelter, A.S. (2024). RNA encodes physical information. bioRxivorg. 10.1101/2024.12.11.627970.

127. Van Treeck, B., and Parker, R. (2018). Emerging roles for intermolecular RNA-RNA interactions in RNP assemblies. Cell 174, 791–802.

128. Luchelli, L., Thomas, M.G., and Boccaccio, G.L. (2015). Synaptic control of mRNA translation by reversible assembly of XRN1 bodies. J. Cell Sci. 128, 1542–1554.

129. Formicola, N., Vijayakumar, J., and Besse, F. (2019). Neuronal ribonucleoprotein granules: Dynamic sensors of localized signals. Traffic 20, 639–649.

130. Mateju, D., Eichenberger, B., Voigt, F., Eglinger, J., Roth, G., and Chao, J.A. (2020). Single-Molecule Imaging Reveals Translation of mRNAs Localized to Stress Granules. Cell 183, 1801–1812.e13.

131. Ramat, A., Haidar, A., Garret, C., and Simonelig, M. (2024). Spatial organization of translation and translational repression in two phases of germ granules. Nat Commun 15, 8020.

132. Chen, R., Stainier, W., Dufourt, J., Lagha, M., and Lehmann, R. (2024). Direct observation of translational activation by a ribonucleoprotein granule. Nat Cell Biol 26, 1322–1335.

133. Smith, J., and Bartel, D.P. (2024). The G3BP stress-granule proteins reinforce the translation program of the integrated stress response. bioRxiv. 10.1101/2024.10.04.616305.

134. Mallik, S., Venezian, J., Lobov, A., Heidenreich, M., Garcia-Seisdedos, H., Yeates, T.O., Shiber, A., and Levy, E.D. (2024). Structural determinants of co-translational protein complex assembly. Cell. 10.1016/j.cell.2024.11.013.

135. Collart, M.A., and Weiss, B. (2020). Ribosome pausing, a dangerous necessity for co-translational events. Nucleic Acids Res. 48, 1043–1055.

136. Popper, B., Bürkle, M., Ciccopiedi, G., Marchioretto, M., Forné, I., Imhof, A., Straub, T., Viero, G., Götz, M., and Schieweck, R. (2024). Ribosome inactivation regulates translation elongation in neurons. J. Biol. Chem. 300, 105648.

137. Shah, S., Molinaro, G., Liu, B., Wang, R., Huber, K.M., and Richter, J.D. (2020). FMRP control of ribosome translocation promotes chromatin modifications and alternative splicing of neuronal genes linked to autism. Cell Rep. 30, 4459–4472.e6.

138. Anadolu, M.N., Sun, J., Kailasam, S., Chalkiadaki, K., Krimbacher, K., Li, J.T.-Y., Markova, T., Jafarnejad, S.M., Lefebvre, F., Ortega, J., et al. (2023). Ribosomes in RNA granules are stalled on mRNA sequences that are consensus sites for FMRP association. J. Neurosci. 43, 2440–2459.

139. Kishi, J.Y., Lapan, S.W., Beliveau, B.J., West, E.R., Zhu, A., Sasaki, H.M., Saka, S.K., Wang, Y., Cepko, C.L., and Yin, P. (2019). SABER amplifies FISH: enhanced multiplexed imaging of RNA and DNA in cells and tissues. Nat. Methods 16, 533–544.

140. Hershberg, E.A., Camplisson, C.K., Close, J.L., Attar, S., Chern, R., Liu, Y., Akilesh, S., Nicovich, P.R., and Beliveau, B.J. (2021). PaintSHOP enables the interactive design of transcriptome- and genome-scale oligonucleotide FISH experiments. Nat. Methods 18, 937–944.

141. Banerjee, A.K., Blanco, M.R., Bruce, E.A., Honson, D.D., Chen, L.M., Chow, A., Bhat, P., Ollikainen, N., Quinodoz, S.A., Loney, C., et al. (2020). SARS-CoV-2 disrupts splicing, translation, and protein trafficking to suppress host defenses. Cell 183, 1325–1339.e21.

142. Yu, G., Wang, L.-G., Han, Y., and He, Q.-Y. (2012). clusterProfiler: an R package for comparing biological themes among gene clusters. OMICS 16, 284–287.

143. Kelley, D.R., Snoek, J., and Rinn, J.L. (2016). Basset: learning the regulatory code of the accessible genome with deep convolutional neural networks. Genome Res. 26, 990–999.

144. Zhou, J., and Troyanskaya, O.G. (2015). Predicting effects of noncoding variants with deep learning-based sequence model. Nat. Methods 12, 931–934.

145. Avsec, Ž., Weilert, M., Shrikumar, A., Krueger, S., Alexandari, A., Dalal, K., Fropf, R., McAnany, C., Gagneur, J., Kundaje, A., et al. (2021). Base-resolution models of transcription-factor binding reveal soft motif syntax. Nat. Genet. 53, 354–366.

146. Uhl, M., Tran, V.D., Heyl, F., and Backofen, R. (2021). RNAProt: an efficient and feature-rich RNA binding protein binding site predictor. Gigascience 10, 1–13.

147. Quang, D., and Xie, X. (2016). DanQ: a hybrid convolutional and recurrent deep neural network for quantifying the function of DNA sequences. Nucleic Acids Res. 44, e107–e107.

148. Avsec, Ž., Agarwal, V., Visentin, D., Ledsam, J.R., Grabska-Barwinska, A., Taylor, K.R., Assael, Y., Jumper, J., Kohli, P., and Kelley, D.R. (2021). Effective gene expression prediction from sequence by integrating long-range interactions. Nat. Methods 18, 1196–1203.

149. Kingma, D.P., and Ba, J. (2014). Adam: A method for stochastic optimization. arXiv [cs.LG].

150. Liaw, R., Liang, E., Nishihara, R., Moritz, P., Gonzalez, J.E., and Stoica, I. (2018). Tune: A research platform for distributed model selection and training. arXiv [cs.LG].

